# The medial entorhinal cortex is necessary for the stimulus control over hippocampal place fields by distal, but not proximal, landmarks

**DOI:** 10.1101/2022.08.04.502628

**Authors:** Elizabeth A.M.A. Allison, Joe W. Moore, Daisy Arkell, Paul A. Dudchenko, Emma R. Wood

**Author notes:** Correspondence should be addressed to Emma Wood at. The authors would like to thank Ms Julia Thomas for her assistance with this study. Elizabeth Allison and Joe Moore are the joint first authors of this work.

## Abstract

A fundamental property of place cells in the hippocampus is the anchoring of their firing fields to salient landmarks within the environment. However, it is unclear how such information reaches the hippocampus. In the current experiment, we tested the hypothesis that the stimulus control exerted by distal visual landmarks requires input from the medial entorhinal cortex (MEC). Place cells were recorded from mice with ibotenic acid lesions of the MEC (n = 7) and from sham-lesioned mice (n = 6) following 90° rotations of either distal landmarks or proximal cues in a cue controlled environment. We found that lesions of the MEC impaired the anchoring of place fields to distal landmarks, but not proximal cues. We also observed that, relative to sham-lesioned mice, place cells in animals with MEC lesions exhibited significantly reduced spatial information and increased sparsity. These results support the view that distal landmark information reaches the hippocampus via the MEC, but that proximal cue information can do so via an alternative neural pathway.

## Introduction

A fundamental property of hippocampal place cells is that their place fields are typically anchored to salient landmarks in the recording environment. For this to happen, landmark information must reach the hippocampus and an association between place fields and the landmark(s) must occur. The traditional demonstration of this association is the cue card rotation manipulation developed by Muller and Kubie (1987). They showed that in a cylindrical recording chamber with a sole polarising visual landmark - a cue card affixed to the chamber wall - rotation of the cue card was associated with a comparable rotation in the location of place fields. This stimulus control by salient landmarks has also been shown in head direction, border and grid cells (Taube et al., 1990; Solstad et al., 2008; Sargolini et al., 2006), and also extends to spatial behavior (Suzuki et al., 1980; Dudchenko et al., 1997).

How does landmark information access place cells in the hippocampus? A likely cortical input for such information is the entorhinal cortex (EC), itself a convergence site from other cortical regions. The EC is divided into two regions, the medial entorhinal cortex (MEC) and the lateral entorhinal cortex (LEC). Electrophysiological recordings from the MEC have revealed several different spatially modulated cell types, with neurons encoding an animal’s speed, head direction, and location (grid cells and boundary cells) in an allocentric spatial framework (Taube 2007, Solstad et al., 2008, Fhyn et al., 2004; Kropff et al., 2015), and projections from the MEC to the hippocampus contain these different spatial signals (Zhang et al., 2013, Sun et al., 2015). In contrast, LEC neuronal activity appears to be related to the locations of objects and other local cues within the environment, with recent data indicating that this coding may be within an egocentric framework (Hargreaves et al., 2005; Deshmuck & Knierim, 2011; Tsao et al., 2013; Wang et al., 2018). This had led to the proposal that the MEC provides global spatial information related to distal landmark (as well as self motion information), while the LEC provides local cue and contextual information (Heys et al., 2020; Kerr et al., 2007; Knierim et al., 2014; but also see Save & Sargolini, 2017).

This hypothesis is supported by several findings. First, Neunuebel et al. (2013) showed that, in a double rotation manipultion in which the local cues on an annular track were rotated in one direction and distal cues on the walls in the opposite direction, spatially-tuned MEC neurons rotated with the global reference frame provided by the distal landmarks, whereas LEC neurons activity rotated with the local reference frame provided by the floor textures. Similarly, Savelli et al. (2017) showed that at least a subset of grid cells in the MEC were anchored to distal room cues when the local reference frame (the recording platform) was rotated. More recently, Fernandez-Ruiz et al. (2021) showed that optogenetic disruption at gamma rhythm of the MEC selectively impaired learning of a spatial task, whereas such disruption of the LEC impaired acquisition of an object discrimination task. MEC and LEC disruption also resulted in disruption of place- and object-specific firing in CA3/dentate gyrus cells, respectively.

Behavioural results also implicate the MEC in the processing of distal visual landmark information. Hales et al. (2014) trained rats for 6 days with one configuration of the watermaze, and then changed all the cues as well as the geometry of the room for new learning. Rats with MEC lesions were impaired at learning this new location and showed perseveration in swimming to the old location relative to control animals. Since all the distal cues, including the room geometry, were different, this finding implies that MEC-lesioned rats did not use distal cues in their initial learning, and relied on whatever minimal local cues were present within the watermaze. Control animals, in contrast, did not return to the formerly correct location, suggesting that their initial spatial learning was based on the distal cues in the room. Similarly, Poitreau et al. (2021) found that MEC-lesioned rats were impaired at using a constellation of distal landmarks to identify the precise location of a hidden platform in a water maze task. These animals did show evidence of some spatial learning in a local cue version of the task (with cues within the water maze), though their performance was still impaired relative to control and LEC-lesioned animals. Yoo and Lee (2017) found that inactivation of the MEC, but not the LEC, impaired the use of visual scenes to guide spatial choices on a T-maze. Surprisingly, LEC inactivations impaired the use of visual scenes to guide a non-spatial choice task. Kuruvilla and Ainge (2017) found that rats with LEC lesions were impaired at relearning a spatial task following surgery based on local cues compared to animals with MEC lesions. In this study, however, no difference was observed between MEC- and LEC-lesioned animals in reacquiring a spatial task based on distal landmarks.

On balance, these data led us to hypothesize that distal cue control over hippocampal place cells requires inputs from MEC, whereas local cue control over place cells requires inputs from LEC. While this specific hypothesis has not previously been tested directly, Miller and Best (1980) showed that place cells in rats with large lesions of the EC failed to anchor to distal visual landmarks in the recording environment (a radial maze) whereas those of control animals did so. Instead, the place cells of lesioned animals appeared to be controlled by intramaze cues. However, as the lesions in this study included both the MEC and LEC, it is unclear whether the observed deficits were specifically related to removal of the MEC, the LEC, or both regions.

To provide a direct test of the hypothesis that the MEC is required for the control of hippocampal place cells by distal landmarks, we rotated distal visual landmarks by 90° and found that, in mice with damage to the MEC, these landmarks failed to exert control over the place fields of hippocampal place cells. In contrast, mice with MEC lesions exhibited intact place field rotation when proximal cues (objects) within the recording environment were rotated by 90°. These findings are consistent with the hypothesis that the MEC is essential for providing distal landmark but not proximal cue information to hippocampal place cells.

## Methods

### Subjects

Experimentation was carried out under a UK Home Office project licence, approved by the Animal Welfare and Ethical Review Board (AWERB) of the University of Edinburgh College of Medicine and Veterinary Medicine, and conformed with the UK Animals (Scientific Procedures) Act 1986.

Subjects were 13 male C57/BL6 mice aged around 8 weeks at the start of the experiment. Following surgery, mice were housed individually on a 12hr light/dark cycle and had free access to food and water in the home cage. All recordings occurred during the light phase of the cycle.

### Electrodes

Recording tetrodes were constructed from 17 µm HML-coated platinum(90%)-iridium(10%) wire (California Fine Wire, CA). Four lengths of wire were twisted and heat-annealed together using a heat gun at 240°C for ∼ 6 seconds to form each tetrode. Two tetrodes were then loaded together into each microdrive (Axona Ltd, UK).

Shortly before surgery, the tetrodes were trimmed using ceramic scissors (Fine Science Tools, Germany) under a microscope to ∼ 2 mm longer than the base of the drive. Each tetrode was then plated with gold solution (Neuralynx, MT) and a protective 18 gauge outer cannula was then placed around the drive cannula and held in place with sterile Vaseline.

### Surgical Procedures

Of the 13 mice used in this experiment, 7 received bilateral MEC lesions and a microdrive implant and 6 received sham lesions and a microdrive implant. The mice were randomly assigned to a group and both the recording sessions and initial clustering analysis were performed with the experimenter blind to the experimental group.

The lesions/sham lesions and electrode implants were performed during the same surgery. Mice were anaesthetized using isofluorane gas (Abbott Laboratories, IL) in oxygen. Analgesia was achieved by subcutaneous administration of small animal Rimadyl (Pfitzer Ltd, UK) at a dose of 0.08ml/kg body weight. A subcutaneous injection of 2.5ml isotonic saline and glucose solution was also administered at this time. The eyes were covered throughout surgery with hydrating eye-gel (Viscotears, TX). The scalp was then shaved and cleaned with antiseptic and the mouse was then fixed into a stereotaxic frame (Kopf, CA) using a bite-bar, nose-cone and two non-traumatic ear-bars. The mouse was placed on a thermostatic heat blanket and covered with a drape.

The skull was exposed via a midline scalp incision and holes were drilled at the injection sites. For the MEC lesion mice, a glass micropipette (Drummond Scientific, PA) was lowered into the brain at an angle of 10°forwards in the anterior-posterior plane (Figure 1A). The injection site was just anterior to the transverse sinus, and between 3.5-3.7 mm lateral to the midline. Injections of either 20nl or 40nl of ibotenic acid (Tocris, UK) (10mg/ml, pH7.4 in PBS) were made at depths of 2.8 mm, 2.3 mm, 1.85 mm and 1.4 mm below dura. After each injection, the pipette was left in place for 5 minutes before being raised to the next injection site. Following injections (or for shams, piercing of the dura only), sterile gelatin sponge (Spongostan Special, Ferrosan A/S, Denmark) soaked in saline, was placed onto the brain surface.

**Figure 1:**
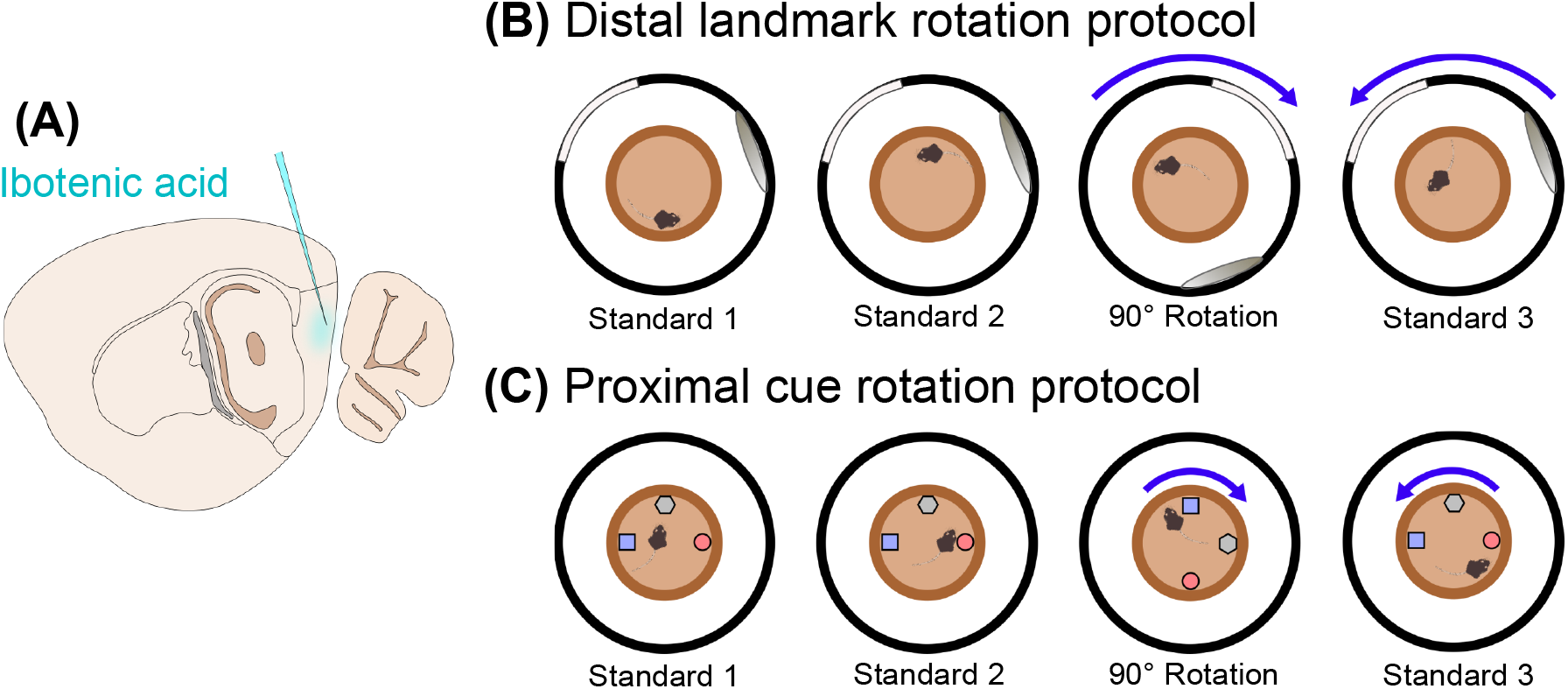
Schematic of injections and landmark rotation protocols **(A)** Schematic of sagittal section of a mouse brain, showing injection of ibotenic acid into the MEC. **(B)** Distal landmark rotation protocol: visual cues were hanging from black curtains surrounding the arena, and mice were given two sessions to explore the central platform with the cues in their standard position, followed by a clockwise rotation of 90°, then a final session with the landmarks returned to the standard position. **(C)** Proximal cue rotation: a similar protocol was used, but with a rotation of objects within the arena, and no cues on the surrounding curtains.

For the microelectrode implant, self-tapping stainless-steel 120TPI screws (Antrin Miniature Specialties Inc, CA) were affixed to the skull and held in place with dental cement (Simplex rapid acrylic denture polymer, Associated Dental Products Ltd, UK). One skull-screw had a grounding wire attached before surgery. The electrode hole was then drilled 2 mm posterior and 2 mm to the right of Bregma and the electrode was lowered into position 0.9 mm below dura (∼ 0.15 mm above the CA1 pyramidal layer). The outer cannula was lowered into position around the electrode above the skull, and sterile Vaseline was used to ensure the join was sealed. The ground wire was soldered onto the skull-screw wire and skull-screws, the base of the drive and the injection sites were all covered over with dental cement. Mice were placed on a heat bench at 30 °C until they fully regained consciousness and then for a further hour of recovery. They were then given 10 days for recovery, during which all mice regained their pre-surgery weight before screening commenced.

### Electrophysiological recording

Mice were connected to a 32-channel recording system (Axona Ltd, UK) via a headstage amplifier and pre-amplifier. Screening occurred while mice explored a cylindrical environment. The signal was amplified, filtered with a bandpass filter at 600-6000 Hz and single-units were identified using the oscilloscope in the DACQ software (Axona Ltd, UK). If suspected neuronal spikes were observed, a trigger was placed at an appropriate amplitude to collect spikes from putative neurons while minimizing collection of noise spikes. A camera placed above the environment and an infra-red LED on the headstage amplifier allowed tracking of the mouse’s position at a frequency of 50 Hz.

### Apparatus

The recording environment was a plastic flowerpot saucer 50 cm across, with a rim 3 cm high and 2 cm wide. This was placed 60 cm off the floor on a stool in the centre of a circular curtained enclosure 2 m in diameter. The curtains were navy blue with 6 possible exits at uniform distances around the enclosure and the ceiling was covered with a white sheet to remove any directional cues. A speaker, lightbulb, camera and recording cable were placed directly above the centre of the environment above the white sheet, with a small hole in the sheet for the camera lens and cable to pass through. During all recordings white noise was played from the speaker to mask any potential directional auditory cues, and all the lights on the outside of the curtains were turned off to reduce any light differences across the environment. Two large distal landmarks were attached to the curtains with safety pins. One was a white sheet which reached from the floor to the ceiling and was 1.4 m wide, the other was a hula-hoop covered with shiny paper to make a circle 1m in diameter and attached so that the base was level with the height of the saucer. These landmarks were attached at an angle of 130° to each other and could also be rotated by 90° clockwise around the enclosure (Figure 1B). For the proximal cue sessions, the distal landmarks were removed and three different local cues (objects) were placed within the flowerpot saucer (Figure 1C). They ranged in height from 6-11 cm and were different in shape, colour and texture. They were placed in three locations to form an isosceles triangle at the edge of the floor of the saucer (as in Save et al., 2005) but as the saucer had a 2 cm rim the mouse could walk around the outside of the objects on top of the rim.

### Rotation sessions

Once multiple place cells had been identified in a screening session, mice were recorded for three days in the environment with distal landmarks followed by three days in the environment with the proximal cues. Two of the control mice and three of the lesion mice were recorded for a further day with the distal landmarks after finishing the local cue recordings, to allow us to test whether any differences between conditions were due to the order in which the conditions occurred. On a given day mice would have 4 or 5 sessions each of 15 minutes with a break of approximately 5 minutes between sessions. For distal landmark and proximal cue rotation days the order of sessions was: Standard 1, Standard 2, rotation 90°clockwise, rotation 90°anticlockwise, Standard 3 (Figure 1B-C).

For each session, the mouse’s home cage was covered with a blanket and carried into the curtained enclosure. The mouse was carried, still covered, between half a turn to two turns around the environment and then removed from the cage, connected to the recording system, and placed in the flowerpot saucer. The mouse foraged for scattered Cheesy Wotsits crumbs (Walkers, UK) for 15 minutes while single-unit and local field potential data were recorded. Following completion of a session, the mouse was unplugged and replaced in its home cage, which was then covered. The saucer was then sprayed and wiped clean with absolute alcohol and placed back in the same orientation (although landmarks moved between sessions the floor of the environment did not). Any necessary rotations were performed during this time. The cage was then picked up and carried round the environment to a random location, before the mouse was taken out, reattached to the recording cable and placed on the recording flowerpot for the next recording session.

### Data analysis

Data files from each session of the day were combined and analysed using a custom Matlab script and a clustering algorithm (KlustaKwik2, developed by Kadir et al., (2014)). Clusters identified by the algorithm were then visualized in Klusters (developed by Hazan et al., (2006)) so that noise clusters could be deleted and incorrect clustering could be fixed. During visual inspection, clusters whose waveforms appeared very similar were combined and irregular spikes judged to be noise were removed from clusters where possible. Custom MATLAB scripts were then used on the outputs to calculate average waveforms for each channel of the tetrode, autocorrelograms, waveform width, isolation distance and L-ratio for each cluster, which were used to visually assess cluster quality. Firing rate maps with 2.5 cm^2^ bins were also generated, and mean firing rate, peak firing rate, waveform width, spatial information content, and sparsity were calculated for each cluster in each session. To calculate in-field mean and peak firing rates, place fields were defined as 9 or more contiguous pixels with firing rates above 20% of the peak firing rate of the rate map, and the mean and peak firing rates were determined from those pixels (Park et al., 2011). The speed of the mouse was calculated for every 500 ms bin, and a threshold of 3 cm/s was applied so that only spikes recorded while the mouse traveled above this speed were included in the analysis.

Cells were identified as pyramidal neurons if they had a mean firing rate between 0.1-5Hz, and waveform width greater than 250 µs, as well as passing visual inspection of waveforms and autocorrelograms. Clusters that did not meet these criteria were excluded from further analyses. Sparsity was calculated using the following equation 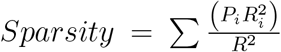 where *i* is the bin number in the firing rate map, *P*_*i*_ is the probability that bin *i* is occupied, *R*_*i*_ is the mean firing rate in bin *i*, and *R* is the overall firing rate. Sparsity measures in what proportion of the environment explored by the mouse did spikes occur. Spatial information content (SI) was calculated using the following equation: 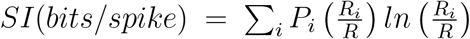, where *i* is the bin number in the firing rate map, *P*_*i*_ is the probability that bin i is occupied, *R*_*i*_is the mean firing rate in bin *i* and *R* is the overall mean firing rate. Spatial information content is a measure of how much information about location is carried by one spike (Skaggs et al., 1993).

Overall mean firing rates, in-field mean and peak firing rates, spatial information and sparsity across sessions were then fitted with generalised linear mixed-models (GLMMs) (lme4 package in RStudio (Bateman et al., 2015)), in order to avoid pseudoreplication by treating cells from the same mouse as independent data points, as well as accounting for the variability in number of cells recorded from each mouse. Mouse and cell identity was included as a random effect, and experimental group (Control, MEC lesion), session (Standard 1, Standard 2, Rotation and Standard 3), and the group x session interaction were included as fixed effects. The data were first modelled with only the interaction term and random effect included, and fit was compared to a model without the interaction term included – the p values reported represent likelihood ratio tests between these models. Post-hoc tests were then used to compare between estimates of fixed effects for the individual sessions and groups; p values from these were Bonferronni-corrected to account for multiple comparisons. As the experiments were repeated on three consecutive days, the data were also modelled with recording day included as a factor, to determine whether this had any effect on the results found (included in Supplementary Material).

In order to analyse landmark-rotation and stability of place fields, the separate firing rate maps for each session were analysed in pairs: Standard 1 vs. Standard 2, Standard 2 vs. Rotation, and Rotation vs. Standard 3. For each pair, the two firing rate maps were overlaid and rotated relative to each other in increments of 5°. A Pearson’s correlation between the two maps was calculated at each angle of rotation, and the maximum and minimum correlations and angles of best correlation were obtained. A cell was only included in this analysis if its firing rate in both of the sessions being compared was greater than 0.1 Hz and cells that did not meet this criterion were categorised separately. A Watson-Williams F-test was used to test whether the mean angle of best correlation was the same between groups. In addition a V-test was used to test whether the circular distribution of angle of best correlation was a uniform distribution or whether it showed a distribution with a mean matching the mean angle expected if the place cells were following the landmarks:0°for the Standard 1-Standard 2 pair, 90°for the Standard 2-Rotation pair, and 270°(−90°) for the Rotation-Standard 3 pair.

The cells were then categorised for each session pair comparison based on their maximum and minimum correlation values, angle of maximum correlation and firing rates in each session, into the following groups: stable, target rotation, off-target rotation, remap, inactive and ambiguous. The criteria were as follows:

- *Stable:* mean firing rate > 0.1 Hz in both sessions, maximum correlation > 0.5, minimum correlation < 0, angle of maximum correlation in the range of 330 to 30°.
- *Target rotation:* mean firing rate > 0.1 Hz in both sessions, maximum correlation > 0.5, minimum correlation < 0, angle of maximum correlation in the range of 240°to 300° for Standard 2 vs rotation (clockwise), 60° to 120° for rotation vs Standard 3 (anticlockwise).
- *Off-target rotation:* mean firing rate > 0.1 Hz in both sessions, maximum correlation > 0.5, minimum correlation < 0, angle of maximum correlation not in the range of the target rotation.
- *Remap:* mean firing rate > 0.1 Hz in both sessions, maximum correlation < 0.5.
- *Inactive:* mean firing rate < 0.1 Hz in either of the two sessions.
- *Ambiguous:* mean firing rate > 0.1 Hz in both sessions, maximum correlation > 0.5, minimum correlation > 0.5 Hz (often because of the field being close to the centre of the arena, so was rotationally symmetrical, so therefore whether it was rotating with the cues cannot be determined).

The proportions of cells which remapped, stayed stable or rotated were calculated for the sham group and the lesion group and a Chi-Square test was used to determine whether the distributions differed. Proportions of cells from each mouse categorised as ‘stable’ and ‘target rotation’ were compared between groups across the sessions, using a two-way ANOVA. In order to compare spatial stability between sessions, correlation values at 0° rotation were compared using a GLMM, as described above.

In case lower precision of place fields in the lesion group skewed results, the rotation analysis was repeated after only including cells with a spatial information content above thresholds of either 0.5bits/spike or 0.3bits/spike. Since many of the cells, particularly in the MEC lesion group had lower spatial information, this greatly reduced the number of cells, but still enabled trends to be observed in the data.

In addition, the analyses of firing rate, spatial information, sparsity and rotation angle were also applied after removing clusters which were identified to be of poor quality. Cluster quality was assessed using a combination of Lratio and isolation distance (ID) (see Schmitzer-Torbert et al., 2005 for equations and validation of the method). Clusters were classed as excellent if ID>30 and Lratio<0.1, good if ID>20 and Lratio<0.15 and acceptable if ID>15 or Lratio<0.2. The values calculated for spatial information, sparsity, firing rate and rotation were recalculated for each tier of cluster quality to identify whether including cells with poor cluster quality had an effect on the results.

### Histology

Following completion of data collection, mice were anaesthetized with isofluorane and given a lethal dose of sodium pentobarbitol (Euthatal, Meridal Animal Health, UK). The tissues were fixed by transcardial perfusion of ice-cold 4% paraformaldehyde. The brains were then extracted and stored overnight at 4 °C in 4% paraformaldehyde before being cryoprotected in 30% sucrose solution. Brains were sectioned in the sagittal plane at 32 µm thickness with a cryostat-microtome. Half of the sections were mounted on polysine slides (Thermo Scientific, UK), stained with 0.1% cresyl-violet, and coverslipped in DPX (Sigma-Aldrich, UK). Sections were then mounted and photographed at 10x magnification using ImagePro Plus (Media Cybernetics, USA). The area of MEC, ventral presubiculum and ventral hippocampus were then calculated for the control animals by drawing around each region on the micrograph, and using ImageJ (NIH, USA) to measure the area. This was then averaged between the control animals. The total area of spared MEC, ventral presubiculum and ventral hippocampus was then calculated for each lesion animal and the percentage of tissue lesioned was calculated. To identify spared tissue, the sections were also examined at 20-30x magnification to determine whether spared regions of tissue contained neurons or only glia. If tissue contained any neurons, it was counted as healthy tissue, but if only glia were present it was counted as scar tissue and was not included in the total area of spared tissue.

## Results

### Infusions of ibotenic acid produced variable amounts of damage to the medial entorhinal cortex

The area of the MEC affected by the lesion varied between the 7 mice in the lesion group. Representative micrographs from a Sham-lesioned brain, a small MEC lesion and a large MEC lesion are shown in Figure 2A-C. For the analyses, all MEC-lesioned animals were considered together, though a breakdown of the average response for each animal and their lesion sizes are presented later in this section. For the mice with ibotenic acid infusions into the MEC, the amount of MEC cell loss ranged from 30% to 94%. For four of these mice, the cell loss was restricted to the MEC. For the remainder, some cell loss was also observed in the ventral hippocampus and the pre-/parasubiculum.

**Figure 2:**
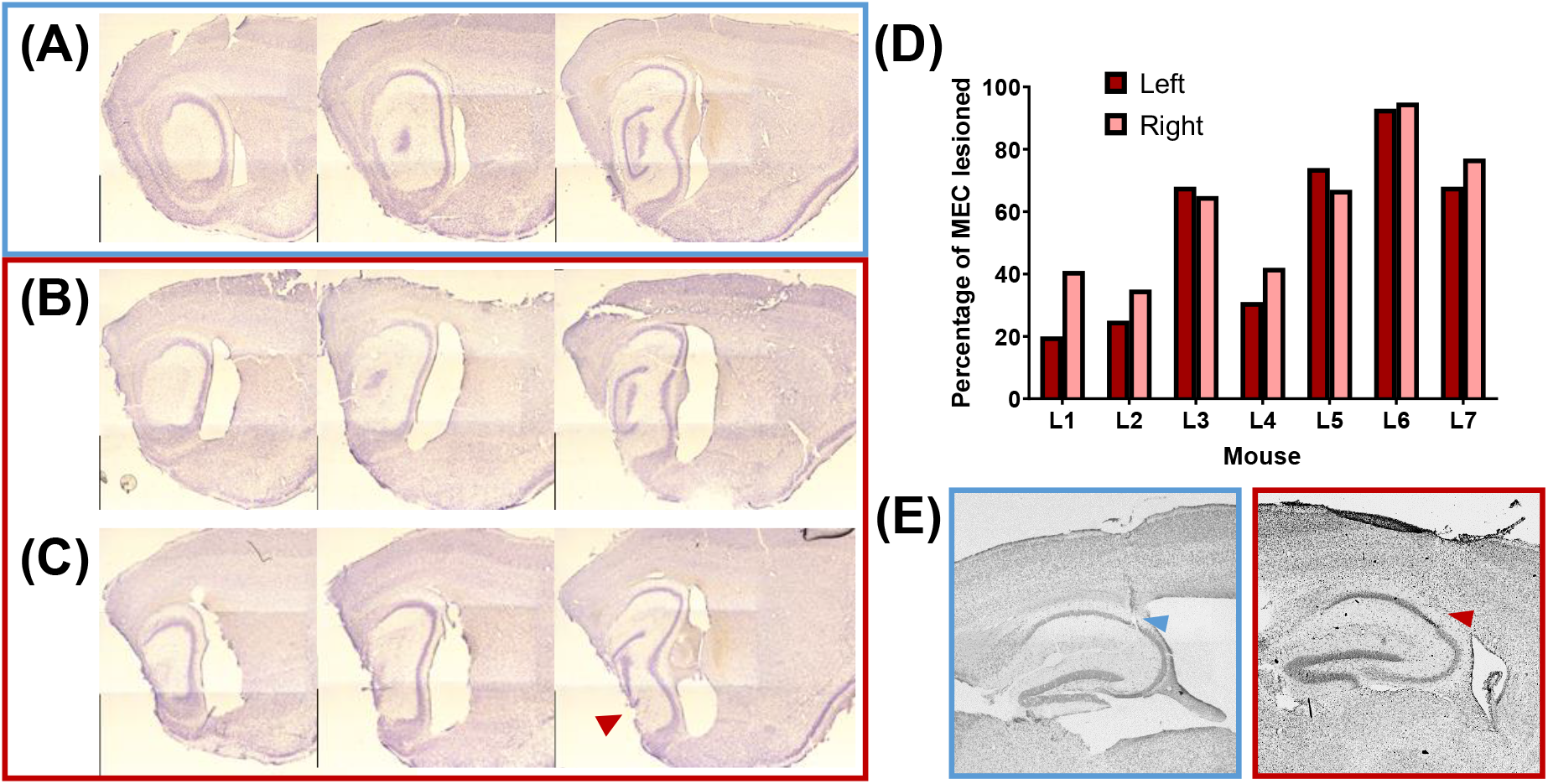
Lesion histology **(A)** Example sagittal sections from a control (sham lesion) mouse brain, showing an intact MEC from lateral (left) to medial (right). **(B)** Similar sections from a lesion mouse with around 70% of the MEC lesioned, with good specificity. **(C)** Example sections from a lesion mouse with a larger, less specific lesion, with almost 100% MEC lesioned, but some damage to other areas, such as ventral hippocampus, as indicated by the red arrow. **(D)** Percentage of the MEC lesioned in the left and right hemispheres of each mouse in the lesion group (n = 7). **(E)** Example electrode tracks for one control mouse (blue box, left) and one MEC lesion mouse (red box, right).

Figure 2E shows two representative examples of electrode tracks, from one control mouse and one MEC lesion mouse. Similar electrode positions in the pyramidal cell layer of dorsal CA1 were observed between mice, with some variability in anteroposterior and mediolateral coordinates (Supplementary Figure 1).

### Place fields in MEC-lesioned animals exhibited lower spatial information and higher sparsity than those in sham-lesioned animals, but no differences in firing rate

In the distal landmark sessions, 376 putative pyramidal cells were recorded from the 6 control mice, and 212 cells were recorded from the 7 lesion mice. Generalised linear mixed-modelling (GLMM) of mean firing rate across the distal landmark sessions, including mice and cells as random factors, found no group x session interaction (likelihood ratio test = 3.33, p = 0.344) and no main effect of group (LRT = 0.409, p = 0.523), but did find a main effect of session (LRT = 43.1, p < 0.0001), with firing rate increasing across the four sessions (Figure 3A). Comparison of within place field mean firing rate found a significant group x session interaction (LRT = 12.8, p = 0.0051), with lesion mice showing generally lower in-field mean firing rates. However, Bonferronni-corrected multiple comparisons between groups for each session found no significant differences (Std1: p = 0.203; Std2: p = 0.823; Rot: p = 0.834; Std3: p > 0.999). Comparison of in-field peak firing rate produced similar results, with a significant interaction (LRT = 11.9, p = 0.0076), but no differences between groups in individual sessions (Std1: p = 0.506; Std2: p > 0.999; Rot: p = 0.931; Std3: p > 0.999) (Figure 3B).

**Figure 3:**
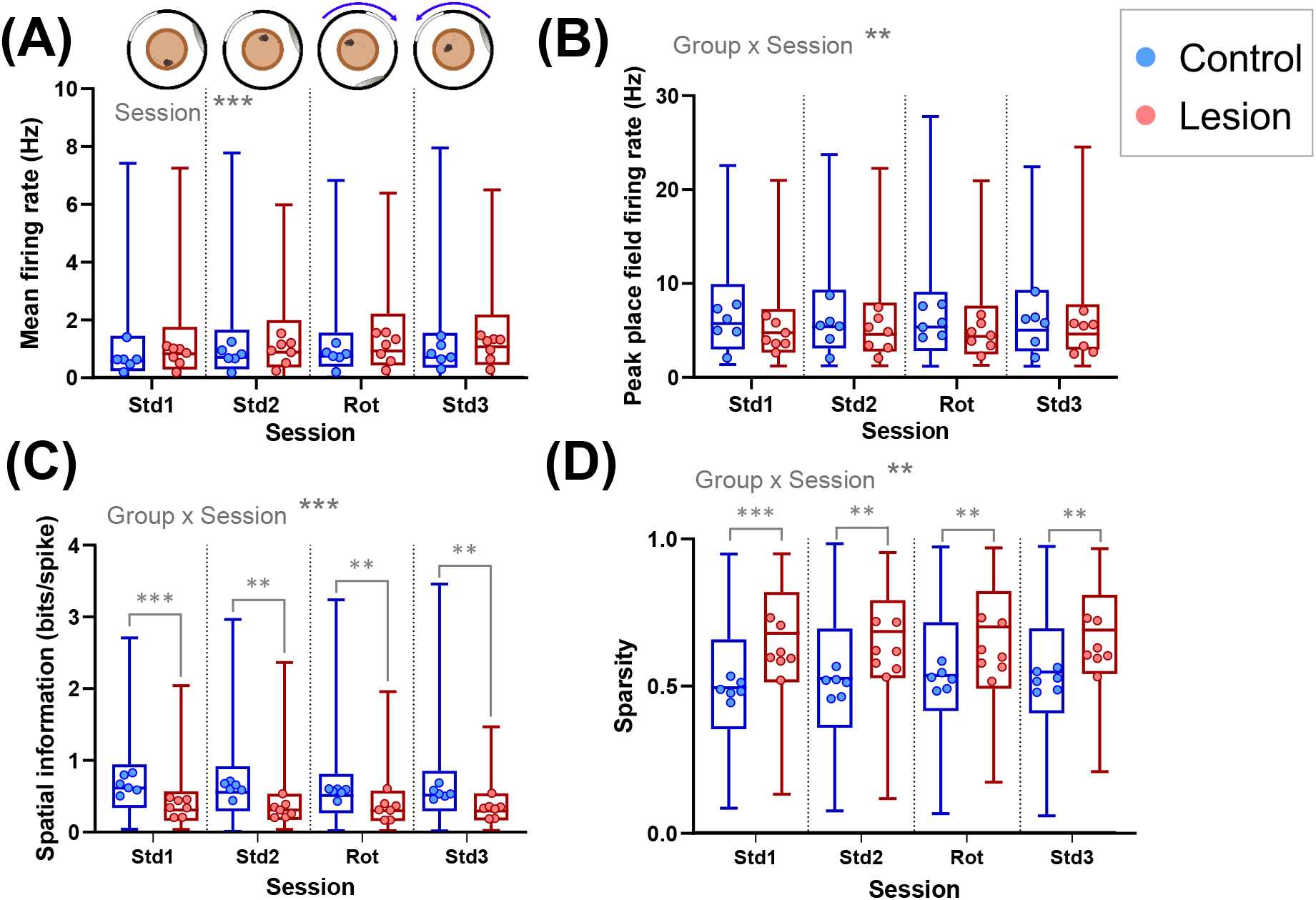
MEC lesion mice place cells have lower spatial selectivity in the distal landmark sessions. **(A)** Mean firing rates for all cells from each group are represented by the box and whisker plots, showing median, interquartile range and range. The mean values for cells recorded from each mouse are shown as dots. **(B)** Peak place field firing rates for all cells from each group are represented by the box and whisker plots, showing median, interquartile range and range. The mean values for cells recorded from each mouse are shown as dots. **(C)** Spatial information values of all active (> 0.1 Hz mean firing rate) cells from each group are represented by the box and whisker plots. The median values for cells recorded from each mouse are shown as dots. **(D)** Sparsity of all active cells from each group are represented by the box and whisker plots. The mean values for cells recorded from each mouse (control n = 6; lesion n = 7) are shown as dots. ** p < 0.01, *** p < 0.001

In contrast, the spatial properties of the cells were affected by MEC lesions. Spatial information was significantly lower in the lesion group, where the median of cells from all mice together was 0.31 bits/spike (interquartile range [0.16,0.57]), while place cells in shamlesioned mice had an overall median of 0.62 bits/spike (IQ range [0.34,0.95]) (Figure 3C). A GLMM of spatial information across the four sessions showed a significant group x session interaction (LRT = 17.5, p = 0.0006), with Bonferronni-corrected post-hoc tests revealing significant differences between the groups in every session (Std1: p < 0.0001; Std2: p =0.0017; Rot: p = 0.0051; Std3: p = 0.0015). The same pattern was seen in the analysis of sparsity, which is another measure of spatial firing; cells from the lesion group fired in a greater proportion of the environment, showing increased sparsity compared to the control group (Figure 3E). A GLMM across sessions showed a significant session x group interaction (LRT = 12.2, p = 0.0067) and revealed significant differences in all four distal landmark sessions (Std1: p = 0.0003; Std2: p = 0.0016; Rot: 0.0051; Std3: p = 0.0024).

The same measures were analysed for the proximal cue sessions. In the proximal cue sessions, fewer cells were recorded 228 pyramidal cells were recorded from the control group, while 178 were recorded from the lesion group. For overall mean firing rate in each session, there was no group x session interaction (LRT = 1.71, p = 0.634), and no main effect of group (LRT = 1.29, p = 0.256), but there was a main effect of session (LRT = 48.0, p = 0.0001) (Figure 4A). However, for in-field mean firing rate, there was a significant group x session interaction (LRT = 23.8, p < 0.0001). In-field mean firing rate was generally lower in the lesion group than the control group, but firing rate for the lesion group increased in later sessions, so post-hoc tests showed significant differences in the first two sessions (Std1: p = 0.020; Std2: 0.012), but no differences in the final two sessions (Rot: p = 0.222; Std3: p > 0.999). In-field peak firing rate showed a similar pattern, with a significant group x session interaction (LRT = 27.5, p < 0.0001), and significant differences between groups in the first two sessions only (Std1: p = 0.033; Std2: p = 0.018; Rot: p = 0.380; Std3: p > 0.999) (Figure 4B).

**Figure 4:**
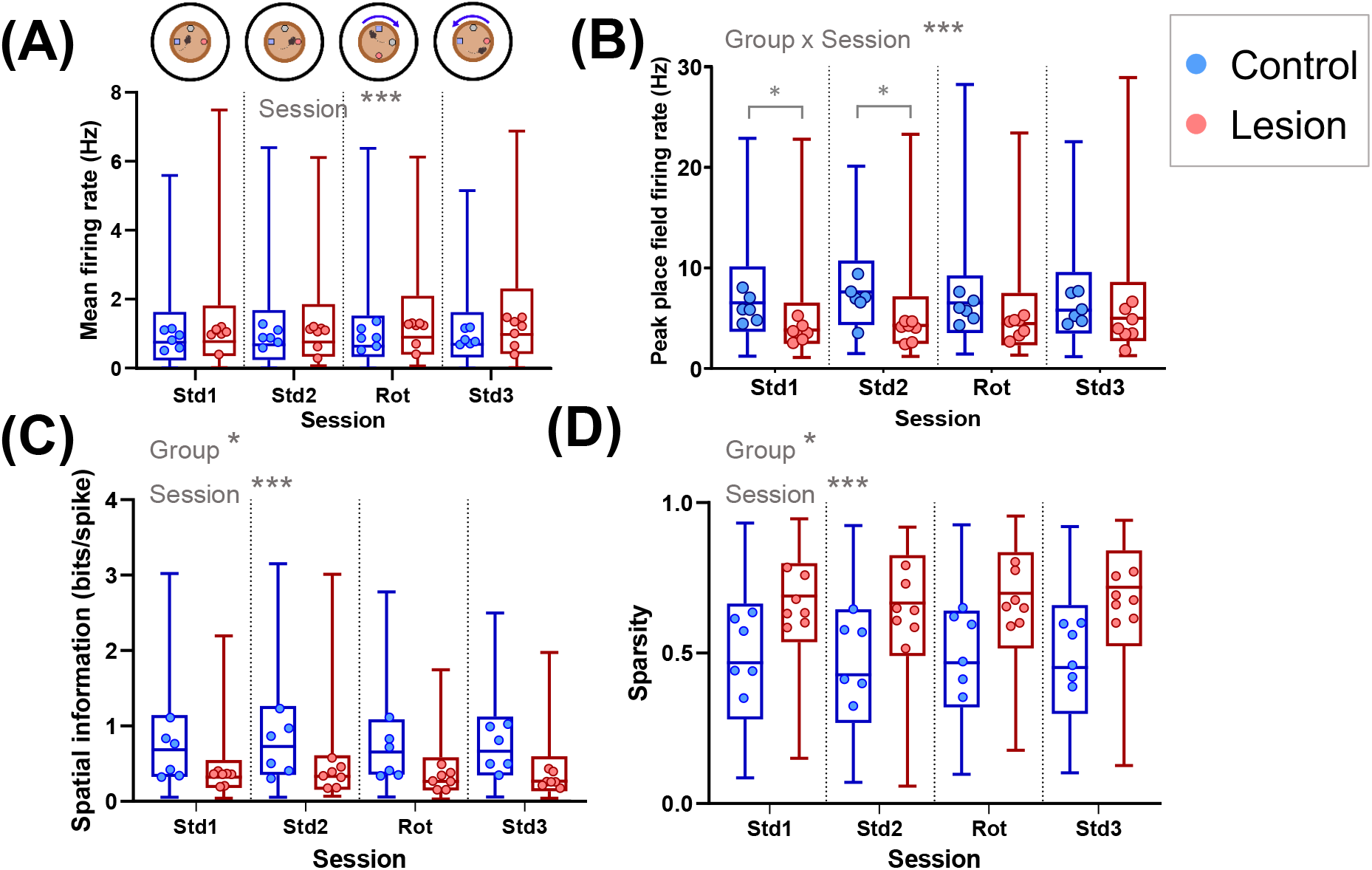
MEC lesion mice place cells have lower spatial selectivity in the proximal cue sessions. **(A)** Mean firing rates for all cells from each group are represented by the box and whisker plots. The mean values for cells recorded from each mouse are shown as dots. **(B)** Peak place field firing rates for all cells from each group are represented by the box and whisker plots, showing median, interquartile range and range. The mean values for cells recorded from each mouse are shown as dots. **(C)** Spatial information values of all active (> 0.1 Hz mean firing rate) cells from each group are represented by the box and whisker plots. The median values for cells recorded from each mouse are shown as dots. **(D)** Sparsity of all active cells from each group are represented by the box and whisker plots. The mean values for cells recorded from each mouse (control n = 6; lesion n = 7) are shown as dots. * p < 0.05, *** p < 0.001

Analysis of spatial properties in the proximal cue sessions also found lower spatial precision. For spatial information, there was no significant group x session interaction (LRT = 5.61, p = 0.132), but there were significant main effects of both group (LRT = 5.79, p = 0.016) and session (LRT = 48.0, p < 0.0001) individually (Figure 4C). Sparsity showed the same effects, with no group x session interaction (LRT = 5.04, p = 0.169), but main effects of both group (LRT = 5.08, p = 0.024) and session (LRT = 40.3, p < 0.0001) (Figure 4D).

These results indicate that the spatial precision of CA1 place cells is reduced in MEC-lesioned mice, shown in both the distal landmark sessions and the proximal cue sessions. The effects of MEC lesions on firing rates are less clear, but in-field firing rates may be reduced slightly in the lesioned mice, although this was only significant in a subset of the proximal cue sessions. As cells were included from repeated recording sessions over three days, the GLMMs were run with recording day included as a factor. This did not alter the conclusions and the results were consistent across the three days (see supplementary material).

### Lesions of the medial entorhinal cortex impair the stimulus control by distal visual landmarks over hippocampal place fields

In the distal landmark rotation condition, place cells from the control group were largely anchored to the distal visual cues and rotated accordingly. In contrast, the place fields of place cells in the lesioned mice were much less likely to follow these cues. This is illustrated in the example place fields shown in Figure 5. The example cells in Figure 5A (blue box) are from control mice, with each row showing the firing rate maps of a cell across the four sessions. The first five rows show cells, each from a different mouse, which rotated their firing fields as predicted by the 90°rotation of the distal landmarks. The bottom row shows an example of a cell that remapped between sessions. The first five rows of Figure 5B (red box) show example cells, each from a different MEC-lesioned mouse, which remained stable across the cue rotation, and the bottom row shows a cell which remapped between sessions.

**Figure 5:**
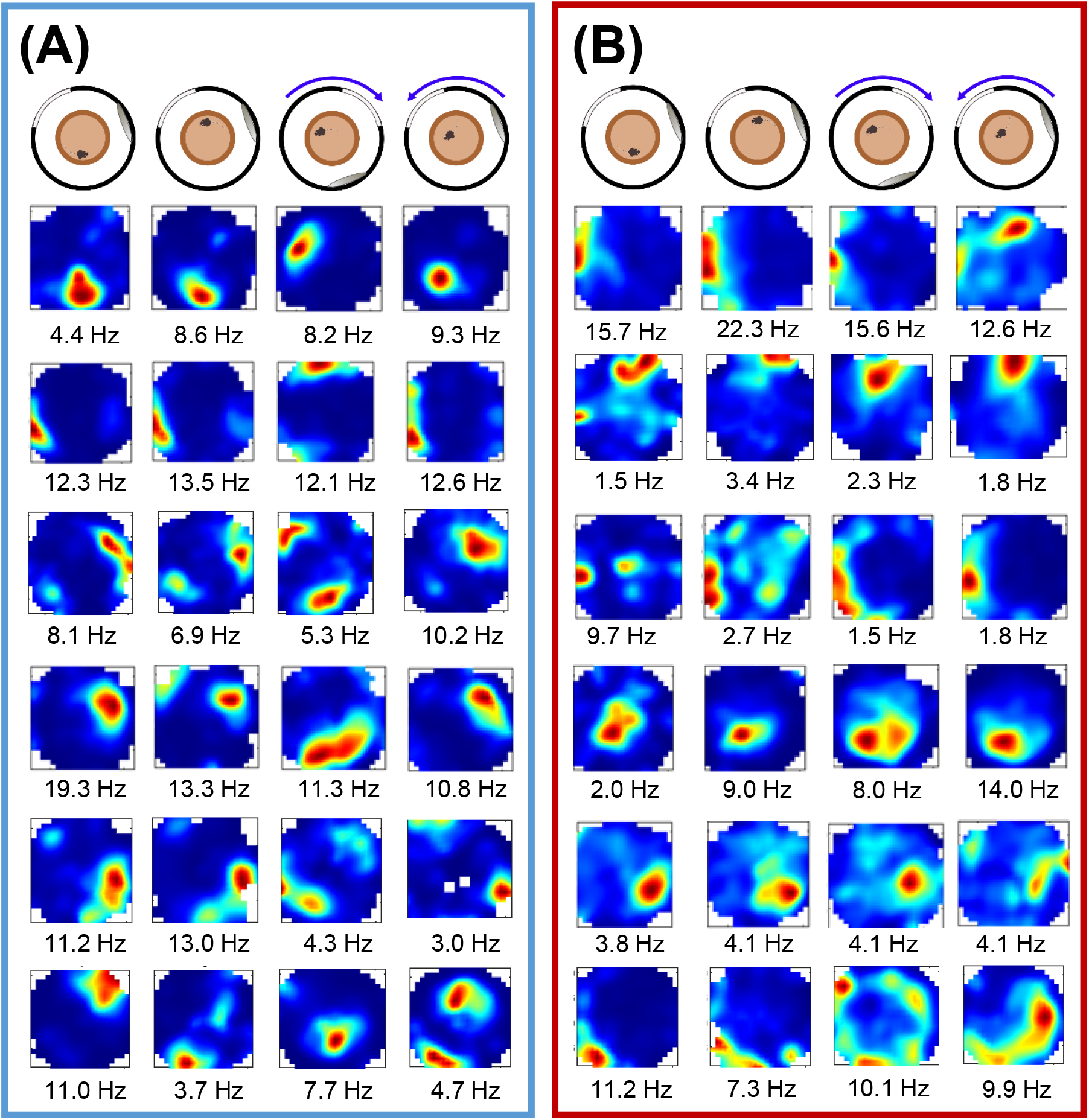
Example place fields across the distal landmark rotation sessions **(A)** Example cells from control mice, showing rotation of spatial firing with the landmarks (top five rows; each cell is from a different mouse), with one example of a remapping cell (bottom row). Peak firing rates are indicated below. **(B)** Place field examples from the MEC-lesioned mice, showing lower spatial selectivity and stability of spatial firing as the landmarks were rotated (top five rows; each cell is from a different mouse).

To quantify the cue rotation data, correlations between firing rate maps were calculated across sessions, rotating one firing rate map relative to the other in 5°increments. The rose plots in Figure 6A show the rotation angles at which the correlation was at its maximum value, for all cells with firing rates of above 0.1 Hz in both sessions being compared, in 10° bins. It is clear that in the sham-lesioned mice (Figure 6A, blue box), cells generally remained stable across the first two standard sessions, when the landmarks did not move, then followed the clockwise 90°rotation of the distal landmarks between the Standard 2 and 90°rotation sessions. The cells then followed the anticlockwise rotation of the landmarks as they were returned to their original position for the final session (Standard 3). Figure 6B also shows that cells from the MEC-lesioned mice were generally stable across the first two standard sessions and remained stable both between Standard 2 and the 90°rotation session (i.e. did not follow the distal landmarks), and when the distal landmarks were returned to their original positions (90°rotation to Standard 3) (Figure 6B, red box). This finding suggests that the distal visual landmarks tended not to exert stimulus control over place fields in the mice with MEC lesions.

**Figure 6:**
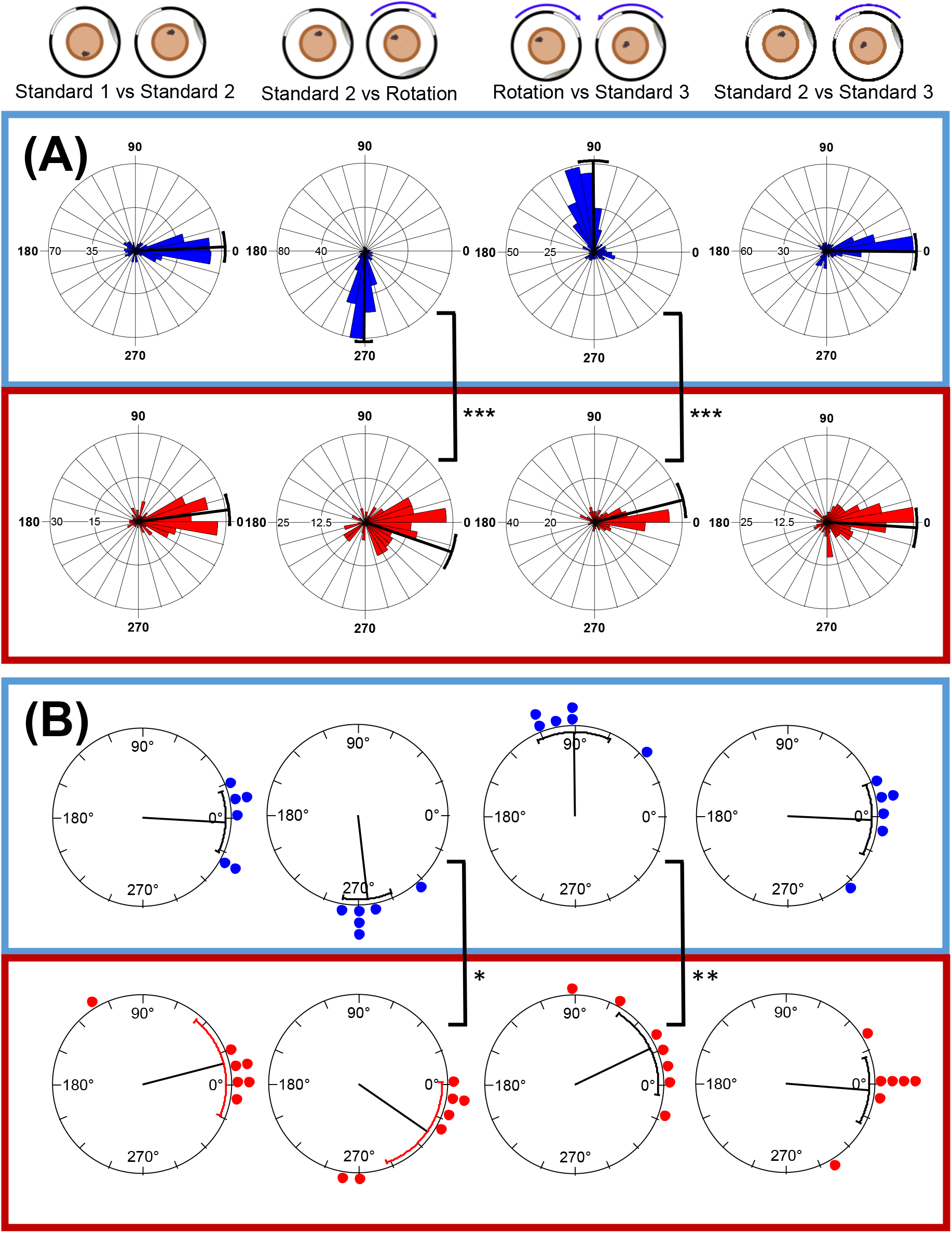
MEC lesion mice place cells fail to rotate with distal landmarks. **(A)** Angular shift at which the maximum correlation between the session rate maps occurred, plotted for cells recorded from all mice in the control (blue) and MEC lesion (red) groups. Radial line denotes circular mean, error bars show 95% confidence intervals. **(B)** Circular mean of maximum correlation angles for cells recorded from each mouse (control n = 6; lesion n = 7). Radial line represents mean of the animal means, with 95% confidence interval error bars. * p < 0.05, ** p < 0.01, *** p < 0.001

Statistical analysis of the distributions of the maximum correlation angles supports this conclusion. When comparing Standard 1 and Standard 2, in both the sham and MEC lesion groups, maximum correlations were clustered around 0°(control: mean vector (µ) of shifts = 1.2°, SE = 5.0°, 95% CI [351.5°, 10.9°]; MEC lesion: µ = 7.4°, SE = 4.8°, 95% CI [358.0°, 16.8°]), suggesting that the place cell firing remained stable across the two sessions. Furthermore, the distributions of the two groups were not significantly different (Watson-Williams F-test: F(1,504) = 0.90, p = 0.34).

Standard 2 and the 90°rotation sessions were then compared; if cells rotated with the distal landmarks they would show maximum correlations around 270°. In the control group, cells were clustered around 270°, (µ = 269.4°, SE = 2.5°, 95% CI [264.3°, 274.3°]) but in the lesion group, there was only a small clockwise shift from 0°in the mean value (µ = 341.0° (−19.0°), SE = 5.5°, 95% CI [330.2°, 351.7°]). This meant that the two distributions were significantly different (Watson-Williams F-test: F(1,531) = 170.4, p < 0.0001).

Comparison of the 90° rotation and the final session, Standard 3, where the landmarks returned to their original position, showed that cells from control mice had maximum correlations clustered around 90°(µ = 90.4°, SE = 4.6°, 95% CI [81.5°, 99.4°]), representing an anticlockwise rotation to follow the cues. Cells from lesion mice again showed only a small rotation (µ = 14.0°, SE = 5.4°, 95% CI [3.4°, 24.5°], and the distributions of rotations in the two groups were again significantly different (F(1,511) = 130.3, p < 0.0001).

As each mouse from each group contributed a different number of cells to the overall total, it is possible that data from mice from which more cells were recorded might bias the data. Therefore, the mean of the maximum correlation values of all cells recorded from each mouse were also compared, and this is shown in Figure 6B. In the control group, for the first two standard sessions, each of the means from the six mice were around 0°, while in the lesion group, most were around 0° with the exception of two mice, and no difference was seen between the groups (F(1,11) = 0.97, p = 0.345). Comparing Standard 2 with the clockwise (−90°) landmark rotation showed that all of the means from the six control mice were clustered around the expected rotation of 270°; in the lesion group, five of the mice appeared to have stable cells, with means close to 0°, while two had means around 270° suggesting that the cells in these two mice rotated with the distal visual landmarks. Despite these two mice that did show rotation with the landmarks, the difference between the groups was significant (F(1,11) = 7.50, p = 0.019). For the anticlockwise rotation back to Standard 3, the mean rotations of the control mice were mainly around 90°as predicted, apart from one mouse showing an under-rotation; the lesion mice showed responses that were consistent with their previous average responses – five remained close to 0°, and the two mice that had rotated with landmarks between Standard 2 and the rotation session showed the expected rotation of around 90°with the landmarks. This difference between the groups was also significant (F(1,11) = 13.89, p = 0.003). Interestingly, the two lesion mice with average responses close to the expected rotations with the landmarks were the two mice with the smallest lesions (see below for further analysis), suggesting the lesions may not have been sufficiently large to cause an impairment.

As an additional measure of cell responses to distal landmark rotations, maximum correlation, minimum correlation, and maximum correlation angles were all taken into account to categorise cells into types of responses. For a cell to be categorised as rotating, as well as having a maximum correlation above 0.5 for a certain angle, it must also have a minimum correlation below 0 for a different angle (see Methods for full classification). In the control group, 56% of cells followed the rotation of the distal landmarks, compared to 4% of the cells recorded in the lesion group. 32% of the cells from the lesion group remained stable when the landmarks were rotated, compared to just 4% in the control group (Figure 7A). The proportions of cells categorised as showing each of the responses listed in Figure 7A were compared across groups. For the first session comparison (Standard 1 vs Standard 2), there was no difference between the groups when using a Chi-square test (*X*^2^(4) = 6.672, p = 0.154). When comparing the sessions across landmark rotations however, there were significant differences in the proportions for both the clockwise rotation (Standard 2 vs Rotation: *X*^2^(5) = 176.9, p < 0.0001) and the anticlockwise rotation back to the standard position (Rotation vs Standard 3: *X*^2^(5) = 128.2, p < 0.0001).

**Figure 7:**
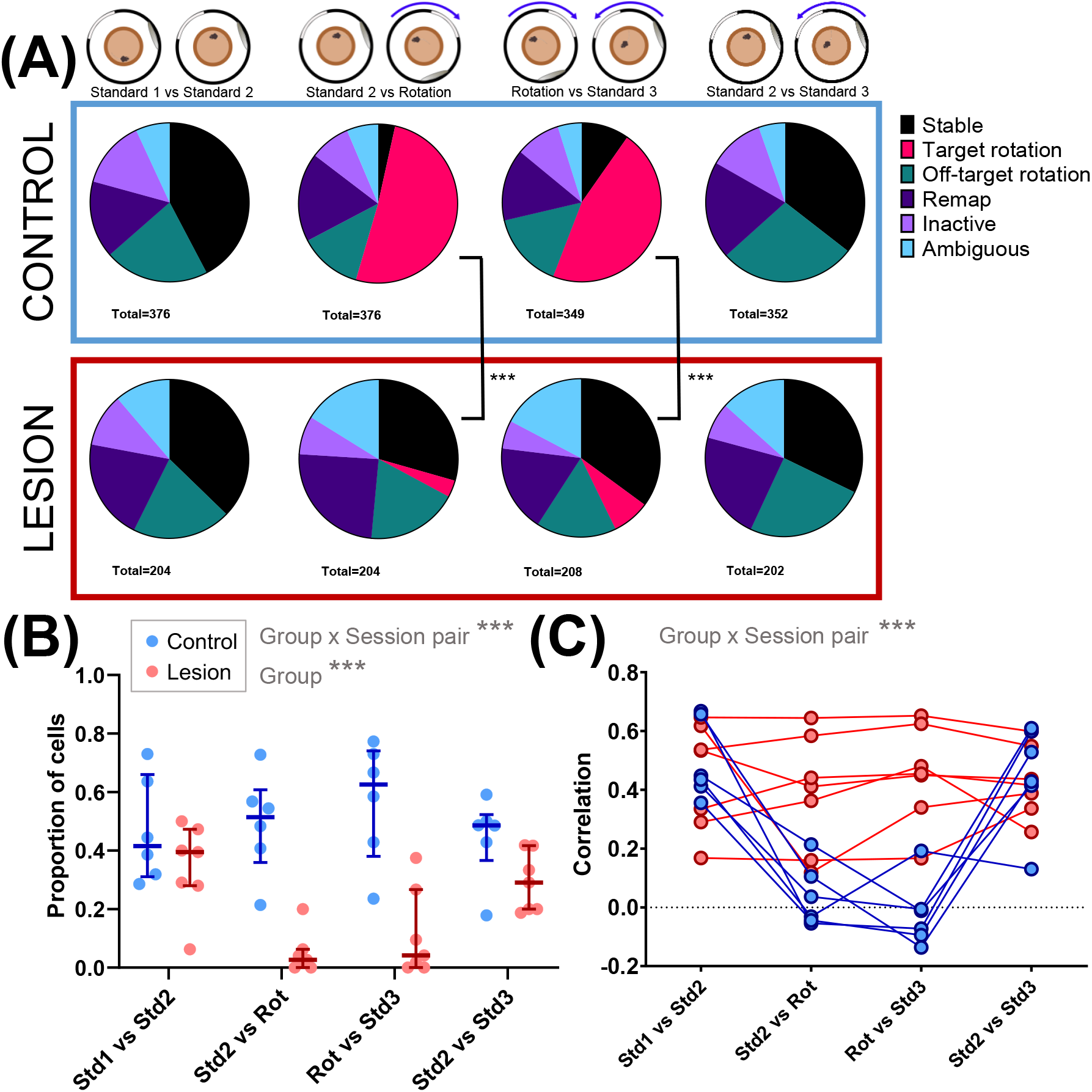
Lower proportions of place cells rotate with the distal landmarks in MEC lesion mice. **(A)** Cells recorded from control (above, blue box) and lesion (below, red box) mice, categorised according to rate map correlations between different sessions, shown as proportions of total number of cells for each session comparison. **(B)** Proportions of cells from each mouse that rotated within ± 30°of the cue rotation angle (Std1 vs Std2: 0°, Std2 vs Rot: 270°, Rot vs Std3: 90°, Std2 vs Std3: 0°) with median ± interquartile ranges plotted (control, blue, n = 6; lesion, red, n = 8). **(C)** Correlation of firing rate maps at 0°, showing means of all active cells recorded from each mouse (control, blue, n = 6; lesion, red, n = 7). Higher correlations indicate more stable firing rate maps. *** p < 0.001

As the analysis described above and displayed in Figure 7A considered all cells from the mice of each group together, and included proportions of cells that showed off-target rotations, remapping, inactivity, or ambiguous responses, the proportions of cells from each individual mouse that either remained stable or rotated within the expected range were compared. Figure 7B compares proportions of cells recorded from each mouse that showed rotation with the cues for each session comparison. Comparison of these values showed a significant main effect of group (F(1,11) = 25.74, p = 0.0004) and a significant interaction between session comparison and group (F(3,33) = 6.95, p = 0.0009). As expected, differences between groups were observed in proportions of cells that rotated with the distal landmarks (Sidak’s multiple comparisons: Standard 2 vs Rotation: p = 0.0034; Rotation vs Standard 3: p = 0.0056), but there were no differences between groups in the proportions that remained stable across the sessions where the cues did not rotate (Standard 1 vs Standard 2: p = 0.608; Standard 2 vs Standard 3: p = 0.200).

The findings described above included data from all three days of the distal landmark condition, raising the possibility that cells inadvertently recorded over more than one day may have biased the findings. The analyses were therefore repeated including only the first day of recording and the same results were observed, even with the reduced number of cells (Supplementary Figure 2). For the subset of mice in which the distal landmark protocol was repeated for a fourth time following the proximal cue sessions, the control mice showed clear rotation of place fields with the distal landmarks, while lesioned mice did not (Supplementary Figure 3).

To determine whether the lower proportion of rotating cells in the MEC lesion group could be due to lower place field stability between sessions, the correlation values at 0°of rotation were compared across the session pairs. Figure 7C shows mean correlation values for cells from each mouse across the session pairs; when comparing sessions across the landmark rotation (Std2 vs Rot and Rot vs Std3), there is a drop in correlation in the control group, as is expected as the cells rotate with the landmarks, so the correlation value at 0°is low. In contrast, the correlations for mice in the lesion group appear to remain at similar values across all session pairs (i.e. both when the landmarks rotated and when they did not). To assess these differences, a GLMM was run on correlations of all active cells (mean firing rate > 0.1) in both sessions in each session pair, with mouse and cell identity as a random factor. A significant group x session pair interaction was observed (LRT: 160.2, p < 0.0001). Post-hoc tests showed that there was no difference between the groups when the cues remained stable, between the first two sessions (Std1 vs Std2: contrast estimate = 0.103, SE = 0.101, p > 0.999), or between Standard 2 and Standard 3 (p > 0.999), suggesting that between-session stability of place cell firing is unaffected by MEC lesion when the distal visual landmarks are stable. As expected, the control group showed a significant drop in correlation in the rotation session pairs, compared to the stable pairs (Std1 vs Std2, compared to Std2 vs Rot: p < 0.0001). However, correlations in the lesion group were not affected by the landmark rotation, and did not differ across the session pairs (Std1 vs Std2 compared to Std2 vs Rot: p > 0.999). Hence there were significant differences between the two groups in stability across the rotation session pairs (Std2 vs Rot: p = 0.0003; Rot vs Std3: p = 0.0002).

These analyses show that the lower proportions of cells rotating with the distal visual landmarks are not likely to be a consequence of reduced overall between-session spatial stability, as similar proportions of cells are stable between the standard (non-rotation) sessions between groups, and this level of stability is maintained across the landmark rotation in the MEC lesion group. It is not known what stable stimuli the MEC lesion mice may use to maintain this stability. The GLMMs were repeated with recording day as a factor and the results showed a similar pattern to when this was not included (see Supplementary Material).

### Lesions of the medial entorhinal cortex do not affect the stimulus control of proximal cues over place fields

In contrast to results from the distal cue rotation, cells recorded in the proximal cue rotations showed more similar responses between the two groups. Figure 8A shows example place cells across the proximal cue sessions for the control mice. The first three rows in Figure 8A show example place cells that showed clear rotation with the proximal objects, while the bottom three show remapping or ambiguous responses across sessions. Figure 8B shows similar examples from MEC lesion mice, with place fields in the first four rows following the 90°rotation with the proximal cues, and the bottom two rows showing less spatially selective cells with remapping or ambiguous responses between sessions.

**Figure 8:**
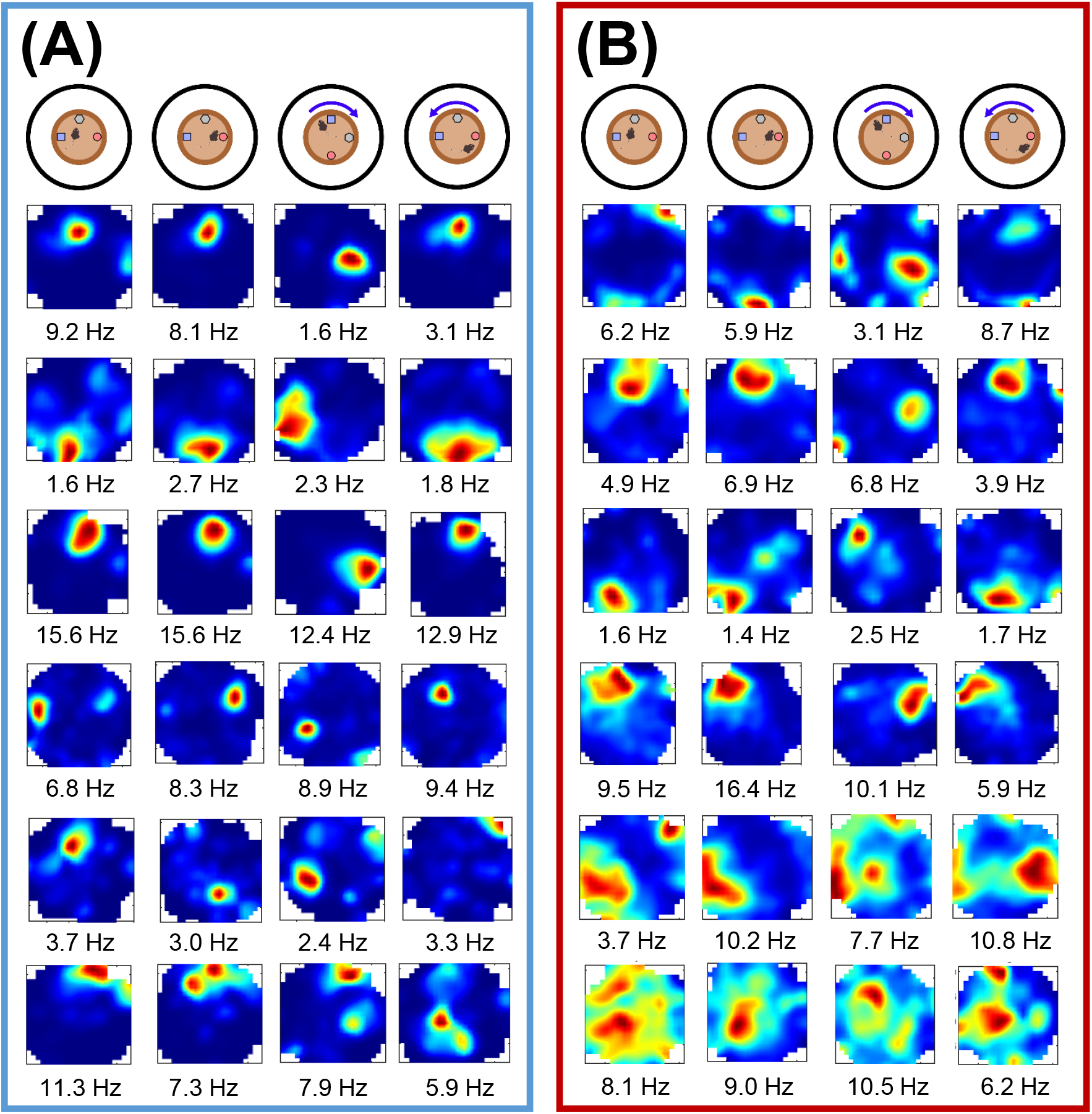
Example place fields across the proximal cue rotation sessions. **(A)** Example cells from control mice, showing cells that rotate with the cues (top three rows), and cells that show remapping across sessions (bottom three rows). Peak firing rates are shown below. **(B)** Examples from the MEC-lesioned mice, showing cells that rotate with the cues (top four rows), and cells with very low spatial selectivity (bottom two rows).

Maximum correlation angles for cells in each group are indicated in Figure 9A. In both groups, maximum correlations between the first two standard sessions were clustered around 0°(control: µ = 2.9°, SE = 4.0°, 95% CI [355.2°, 10.7°]; lesion: µ = 358.3°(−1.7°), SE = 6.2°, 95% CI [346.1°, 10.4°]), suggesting both were remaining stable when the cues were not moved. The two distributions were not significantly different (Watson-Williams F-test: F(1,361) = 0.47, p = 0.49). When comparing Standard 2 to the 90°cue rotation session, the control cells seemed to show a bimodal distribution, with some maximum correlations around the expected rotation of 270°, and some stable at 0°. The mean value was in between these two values at µ = 312.9°(−47.1°) (SE = 7.6°, 95% CI [297.9°, 327.8°]). The cells from the lesion mice appeared to show a slightly more consistent response, generally clustering around 270°, with a mean of µ = 296.7°(−63.3°) (SE = 7.0°, 95% CI [283.0°, 310.4°]). However, the distributions of the correlations of cells in two groups were not significantly different (Watson-Williams F-test: F(1,367) = 3.39, p = 0.07).

**Figure 9:**
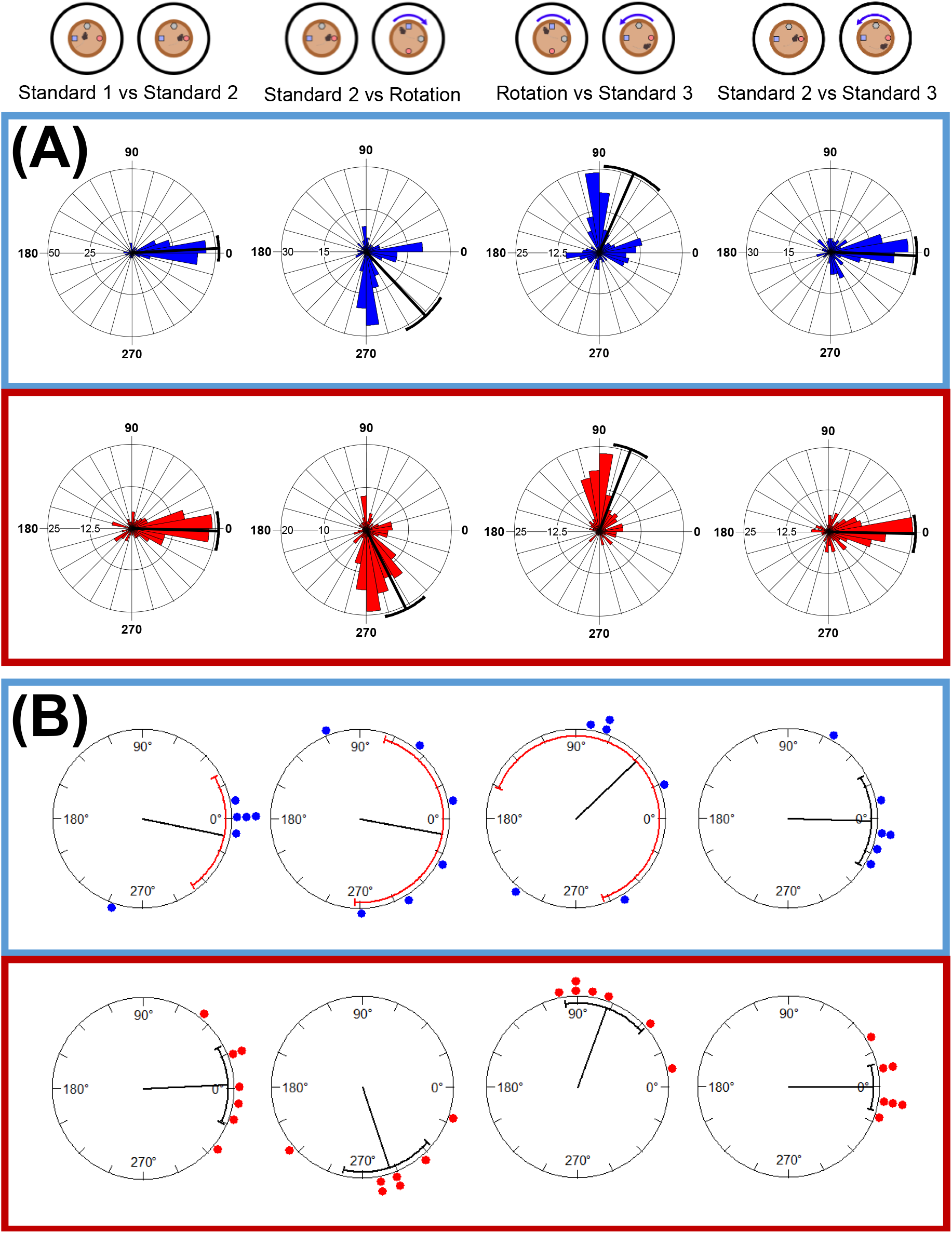
Place cells in both groups rotate with the proximal cues. **(A)** Angular shift at which the maximum correlation between the session rate maps occurred, plotted for cells recorded from all mice in the control (blue) and MEC lesion (red) groups. The radial axis represents number of cells, the bold radial line denotes circular mean, error bars show 95% confidence intervals. **(B)** Circular mean of maximum correlation angles for cells recorded from each mouse (control, blue, n = 6, lesion, red, n = 7). Radial line represents mean of the animal means, with 95% confidence interval error bars.

Finally, comparing the rotation session to Standard 3, most of the control cells were clustered around the expected rotation of 90°, with some remaining stable around 0°(µ = 66.3°,SE = 10.1°, 95% CI [46.6°, 86.1°]), while the lesion cells were again mostly clustered around the expected rotation (µ = 68.7°, SE = 5.7°, 95% CI [57.1°, 79.8°]). Again, there was no significant difference between the two distributions (Watson-Williams F-test: F(1,370) = 0.07, p = 0.79).

Looking at the individual animal mean angle of shift for the proximal cue sessions also suggests the MEC-lesioned mice may have an increased reliance on the proximal cues compared to the controls. The mean rotations for both groups, with the exception of one control mouse, clustered around 0°between the first two standard sessions, and there was no difference between the groups (F(1,11) = 0.384, p = 0.548). The mean rotations of the control mice were much more variable following the 90°rotation of proximal cues than the lesion mice, of which four were close to the expected rotation of 270°(Figure 9B). Because of this high variability, there was again no significant difference between the groups (F(1,11) = 3.00, p = 0.111). When the cues rotated back to the standard configuration, the control mice means were again variable, although three of these means were close to 90°, whereas the lesion mice were again more reliably clustered around the expected rotation, five being close to 90°. Again, there was no significant difference between the groups (F(1,11) = 0.58, p = 0.462). The lower variability in the lesion mice compared to controls (indicated by the radial error bars in Figure 9B) across the rotation sessions suggests a more consistent response of the place fields in the lesion mice. This may indicate that the lack of distal cue information normally conferred by the MEC leaves the place cells of lesioned mice more likely to rely on proximal cue information rather than any uncontrolled distal cues, and therefore to show a more coordinated rotation with the objects.

In both groups, 30% of the cells showed rotations consistent with the 90° cue rotation. However, a slightly higher proportion of cells from the control group (20%) remained stable across the rotation, compared to the lesion group (9%). Chi-square tests found significant differences between the groups in all of the session comparisons (Figure 10A; Standard 1 vs Standard 2: *X*^2^(4) = 35.09, p < 0.0001; Standard 2 vs Rotation: *X*^2^(5) = 25.00, p = 0.0001; Rotation vs Standard 3: *X*^2^(5) = 28.95, p < 0.0001; Standard 2 vs Standard 3: *X*^2^(4) = 12.35, p = 0.0149). Comparing proportions of cells from each mouse that showed rotations following the cues (Figure 10B) showed a significant main effect of session (F(2.61,28.75) = 3.91, p = 0.0226), but no significant effect of group (F(1,11) = 0.04, p = 0.843) and no significant interaction (F(3,33) = 2.88, p = 0.0509). Multiple comparisons did not reveal any significant differences between the groups for any of the comparisons.

**Figure 10:**
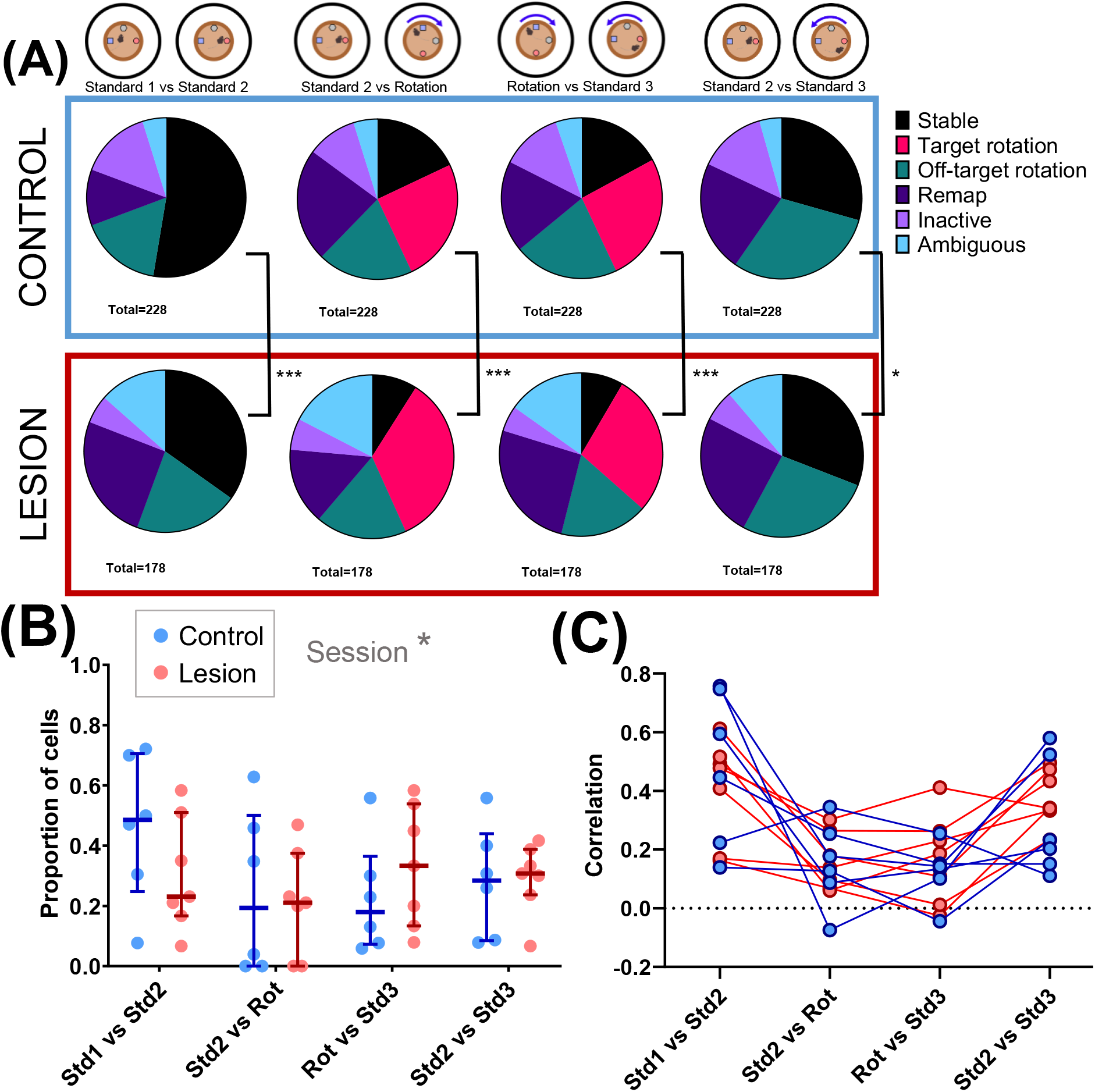
Similar proportions of cells rotate with the proximal cues in both groups. **(A)** Cells recorded from control (above, blue box) and lesion (below, red box) mice, categorised according to rate map correlations between different sessions, shown as proportions of total number of cells for each session comparison. **(B)** Proportions of cells from each mouse that rotated within ± 30°of the expected rotation angle, with median ± interquartile ranges plotted (control, blue, n = 6; lesion, red, n = 7). * p < 0.05 **(C)** Correlation of firing rate maps at 0°, showing means of all active cells recorded from each mouse (control, blue, n = 6; lesion, red, n = 7). * p < 0.05, *** p < 0.0001

Figure 10C shows the mean correlation values at 0° of rotation for each mouse across the different session comparisons. From this, mice from both groups generally show a reduction in mean correlation values for the rotation comparisons compared to the stable comparisons. A GLMM was fitted to these data, which found a significant group x session interaction (LRT = 12.1, p = 0.0071). However, there were no differences in correlation values between groups for any of the session comparisons. There was no difference in stability between the groups when the objects remained stable (Std1 vs Std2: p = 0.270), and there was a similar decrease in correlation at 0°for both groups when the objects were rotated (Std1 vs Std2 compared to Std2 vs Rot: Control: p < 0.0001; Lesion: p < 0.0001), and so there was no difference between the groups comparing the across the cue rotation (Std2 vs Rot: p > 0.999).

### Larger lesions of the MEC were associated with less stimulus control over place fields by distal landmarks

As mentioned in the previous sections, lesion size appeared to have an influence on some of the results observed. All mice with more than 36% of MEC lesioned had average rotations close to 0°when the distal landmarks were rotated (see Supplementary Material). In contrast, the two mice with smaller lesions had average rotations consistent with the 90° rotation, suggesting an association between MEC lesion size and rotation with distal landmarks. Figure 11A shows proportions of cells that were categorised as following the initial clockwise distal landmark rotation (Standard 2 vs Rotation) for each mouse, plotted against percentage of lesioned MEC. A trend can be seen, with larger lesions being associated with lower proportions of rotating cells. This correlation was significant (Spearman’s rank correlation: r(7) = −0.927, p = 0.007). While correlation of proportion of cells rotating with the proximal cues with lesion size did show a positive trend (Figure 11B), this did not reach significance (Spearman’s correlation: r(7) = 0.700, p = 0.090).

**Figure 11:**
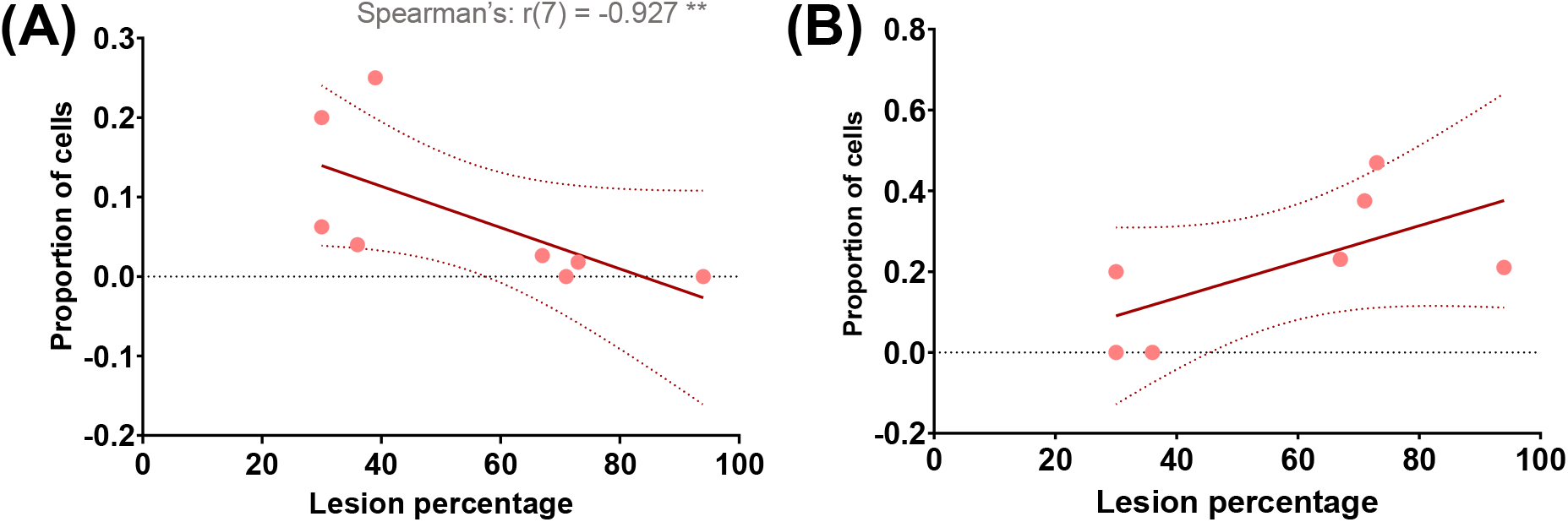
Larger MEC lesions are associated with lower proportions of cells rotating with distal landmarks. **(A)** Proportion of cells recorded from each lesion mouse that rotated within ± 30° of the initial clockwise rotation in the distal cue sessions, against percentage of MEC lesioned. The data were fitted with a linear regression line with dotted lines showing 95% confidence intervals. **(B)** Proportion of cells recorded from each lesion mouse that rotated within ± 30° of the initial clockwise rotation in the proximal cue sessions, against percentage of MEC lesioned. The data were fitted with a linear regression line with dotted lines showing 95% confidence interval.

Finally, we also observed that damage encroachment of the lesions to surrounding areas did not have any systematic effect on rotation with the cues. Animals with large lesions of the MEC (> 60% tissue loss) either with or without extra-MEC damage showed comparable responses to the distal cue rotations (see Supplementary Material).

## Discussion

The current study tested whether the medial entorhinal cortex is an essential node for distal landmark control over hippocampal place cell fields. We observed, first, that pyramidal cells still exhibited spatial firing in the absence of an intact MEC in mice. This is consistent with previous findings in rats (Miller and Best 1980; Brun et al., 2008; Van Cauter et al., 2008; Hales et al., 2014; Ormond and McNaughton 2015; Jacob et al., 2020). Second, and the main finding of the current study, our results show that disruption of MEC input to the hippocampus impairs the stimulus control by distal landmarks over hippocampal CA1 place fields. Finally, our results show that stimulus control by local cues is not disrupted in MEC-lesioned animals, suggesting a shift in their use of landmarks relative to intact animals. We consider these results below.

### MEC lesions impair spatial coding of CA1 place cells in mice

Firstly, this study replicates previous findings of preserved spatial firing in hippocampal place cells, but with reduced spatial information and selectivity when input from the MEC is lost (Van Cauter et al., 2008; Hales et al., 2014; Schlesiger et al., 2015; Jacobs et al., 2020). While these previous studies have used rats, the current experiment provides evidence that a similar effect is observed in mice. Firing rates of CA1 pyramidal cells were only slightly reduced in the current study, with in-field firing rate only significantly different in particular sessions. Previous reports of MEC input manipulation are mixed, with some showing large reductions in hippocampal firing rates (Van Cauter et al., 2008; Hales et al., 2014), and some showing minimal changes (Brun et al., 2008; Schlesiger et al., 2015; Kanter et al., 2017; Jacobs et al., 2020).

The between-session stability of spatial firing did not seem affected by the MEC lesions in the current experiment - correlations between firing rate maps for the first two sessions where the cues were stable and showed no differences between groups. This contrast with the results of Hales et al. (2014), who found a deficit in stability between sessions in MEC lesioned rats. However, this deficit worsened as the delay was increased and at the shortest delay of 2 minutes, comparable to the delay used in this experiment, the deficit was at its lowest (similar results were shown in Schlesiger et al. (2018)). The mean correlation values reported by Hales et al. (2014) were also higher than in this study, so this effect could depend on the type of environment used, or familiarity of the environment. There is some evidence to support that this may be a species difference, as mice show lower levels of place field stability than rats when freely exploring an environment, only showing comparable levels when spatial memory is required for a task (Kentros et al., 2004). Therefore, the lack of task demand in the current experiment may have resulted in relatively low place field stability in both groups of mice, and any effects of MEC lesion may not have been observed for this reason. The stability of cells from the lesion group appeared to remain at similar levels of stability across the distal cue rotations, suggesting they were anchoring to other cues, possibly the limited local features of the platform. This could perhaps explain the lower place field precision, as having less reliable landmark information may decrease accuracy of location estimation.

### The medial entorhinal cortex is essential for stimulus control over CA1 place fields by distal landmarks

In control animals, the distal landmarks used in the current experiment exerted strong stimulus control over hippocampal place fields. This was seen not just in the responses of the majority of place cells, but also in the average response of the place cells for each animal (Figure 6). In contrast, for the MEC-lesioned mice, a 90° rotation of the distal landmarks failed to yield a corresponding rotation of place fields. Indeed, the mean shift for the lesioned mice was 19.5°. At the level of individual animal averaged responses, five lesioned animals show little rotation, two showed an average rotation within 30° of 90°, and one showed a large shift of 146.7°. Four of the five of the MEC-lesioned mice that showed little average place field rotation had larger MEC lesions (> 64% tissue loss), whereas the two lesioned mice which showed roughly appropriate rotations had 30% MEC tissue loss. These results suggest that smaller amounts of damage to the MEC may be insufficient to disrupt distal landmark information flow to the hippocampus.

The disruption of stimulus control indicates that distal landmark information reaches the hippocampus via the MEC. The current results are thus consistent with the impairment in stimulus control observed by Miller and Best (1980) on a radial arm maze, although in their study complete EC lesions were used. The current results are also consistent with the impairments in stimulus control over spatial behaviour in animals with MEC lesions (Parron et al., 2004; Hales et al., 2014; Poitreau et al., 2021).

Of the previous experiments, only Van Cauter et al. (2008) performed cue rotation manipulations. Although they saw a decrease in the proportion of cells showing rotation, they did not find the extensive deficit seen here. One possible reason for this is the types of cues they used. Distal landmarks and cue cards can only be perceived from one side whereas local object cues can be seen from multiple angles and, if not placed at the very edge of an environment, can also be circumnavigated. The cues used by Van Cauter et al. were object cues similar to those used in the proximal cue condition here, but these were placed at the very edge of the environment. Since there appears to be a difference between cues which are at the edge (touching the walls) of an environment, and cues which do not touch the edge of the environment (Scaplen et al., 2014) it is not clear whether their cues would be processed by the distal cue pathway or the object cue pathway (or perhaps both). The effect of MEC lesions in their experiment is somewhere in between the results seen here in the distal landmark and local cue conditions, with increased remapping compared to control animals, but still some evidence of anchoring to cues, with 49% of cells in the lesion group showing place fields which rotated with the cues.

A similar argument applies to the results of Jacobs et al. (2020). They found that MEC-lesioned rats showed less place cell stability compared to sham-lesioned animals in a high-walled, cylindrical environment containing three distinct landmarks on the cylinder periphery. These authors also observed that place cells in the MEC-lesioned animals were even more unstable when the place cells were recorded within this environment in the absence of the landmarks in the dark. These results suggest, indirectly, that the three landmarks (which may have been perceived as distal or proximal, as discussed above) may have exerted some stimulus control over the place fields in the MEC-lesioned rats.

### The medial entorhinal cortex is not essential for stimulus control over place fields by proximal cues

The place fields of mice with MEC lesions tended to rotate with the 90° rotation of the proximal cues, though with some undershoot. This was seen both at the level of individual place fields and the averaged response for six of the seven lesioned animals (with the remaining animal overshooting the rotation by ∼ 50°). The pattern of responses in the sham-lesioned mice was more complicated. There was clear evidence of local cue control with a large number of fields, but also a clear suggestion of a lack of cue control for other fields. At the level of average responses, likewise, 90°cue rotations and return-from-rotation sessions featured variable responses - much more so than in the MEC-lesioned animals (Figure 9). This pattern of results suggests that the MEC is not essential for local cue control over place fields. It also hints at the possibility that, for animals with an intact MEC, local cues (objects) within the environment do not necessarily provide a strong anchoring landmark.

This pattern of results is consistent with the findings from a previous recording study by Neunuebel et al. (2013). They observed strong distal landmark control over a large subset of MEC neurons in a double-rotation manipulation. Their results for LEC neurons, in contrast, were more variable, but were more consistently associated with local cue control. This parcellation of function suggests that the MEC and LEC may provide distal landmark and local cue inputs to the hippocampus (Knierim et al., 2014). As such, damage of the MEC would be expected to impair distal landmark control over place fields, while potentially sparing local cue control - just as we have observed. This may also account, indirectly, for the somewhat surprising observation of stability between standard sessions in the distal landmark condition for the MEC-lesioned mice. If distal landmarks do not anchor place fields in these animals (as the lack of rotations with the 90°landmark rotations suggest), then one might expect variability in place field locations across standard sessions. As this was not observed, the possibility remains that uncontrolled aspects of the local environment (the recording dish) may have served as an orientation cue for the MEC-lesioned mice (Zhang & Manahan-Vaughan, 2013). To address this, it would be of interest to repeat these manipulations in animals with LEC lesions to see if the converse set of findings (intact distal landmark control, impaired local cue control) would be observed.

## Summary

The current study makes three contributions. First, it confirms that the medial entorhinal cortex is not essential for recognisable place cell firing fields. Second, it shows that the medial entorhinal cortex is essential for the normal stimulus control of distal visual landmarks over place fields. Third, it provides evidence that the medial entorhinal cortex is not needed for local landmark control over place fields. This pattern of results is consistent with the conceptualisation of the entorhinal cortex provided by Knierim et al. (2014), where distal landmark information is processed by the medial entorhinal cortex, whereas local landmark information is processed by the lateral entorhinal cortex.

## Data Availability Statement

The data that support the findings of this study are available from the corresponding author upon reasonable request.

## Notes

This research was supported by a grant to P.A.D. and E.R.W. from the Biotechnology and Biological Sciences Research Council (BB/P002455/1).

### Competing Interest Statement

The authors have declared no competing interest.

### Summary of Updates

Author added - forgotten in previous version.

