## Supporting Information for "The medial entorhinal cortex is necessary for the stimulus control over hippocampal place fields by distal, but not proximal, landmarks"

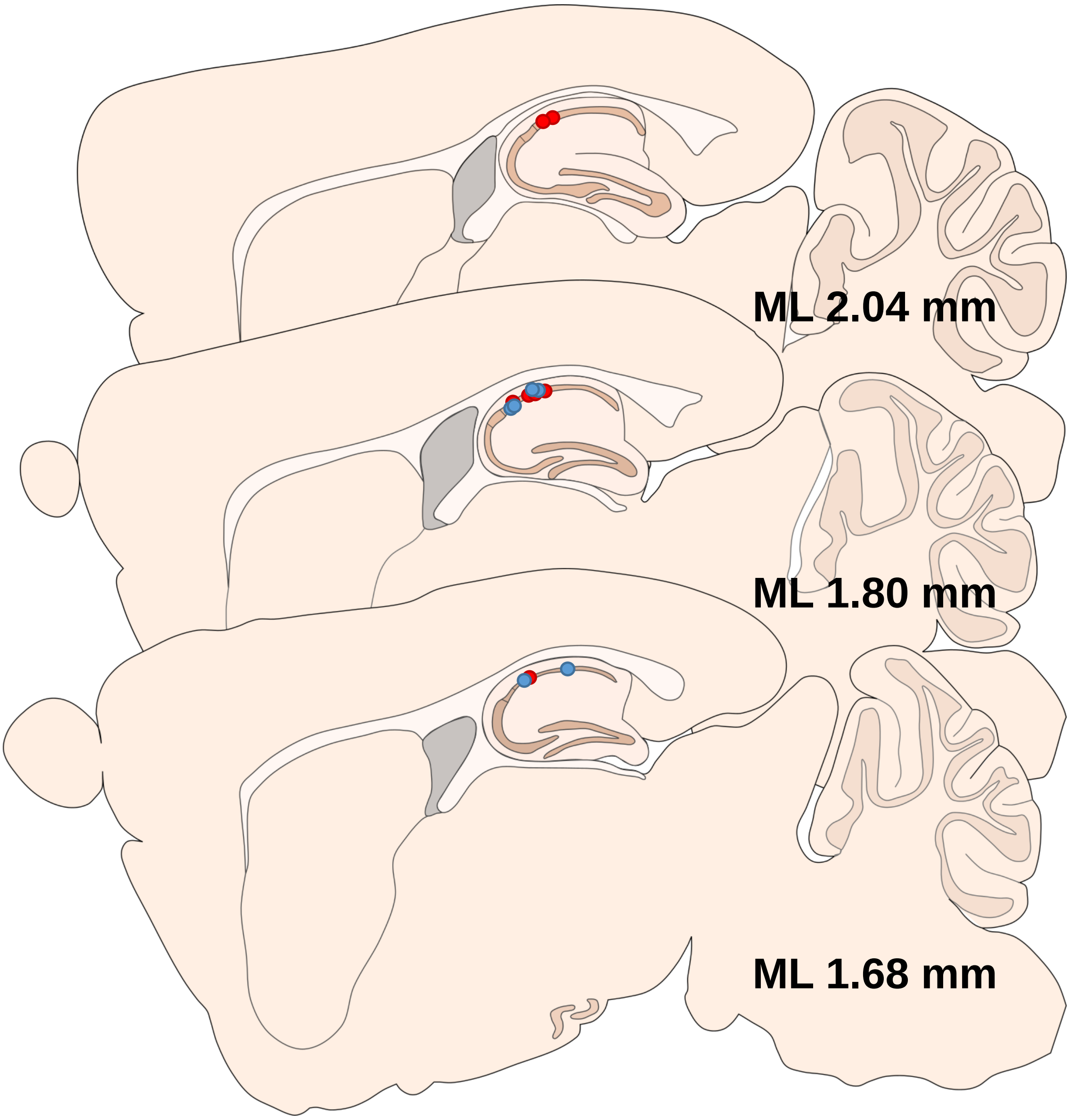


**Figure 1:** Approximate tetrode tip positions of all mice. Blue dots represent control mice (n = 6), red dots represent MEC lesion mice (n = 7). Approximate ML measurements were determined from sagittal mouse brain atlas images that most closely matched the section showing the tetrode tracks.

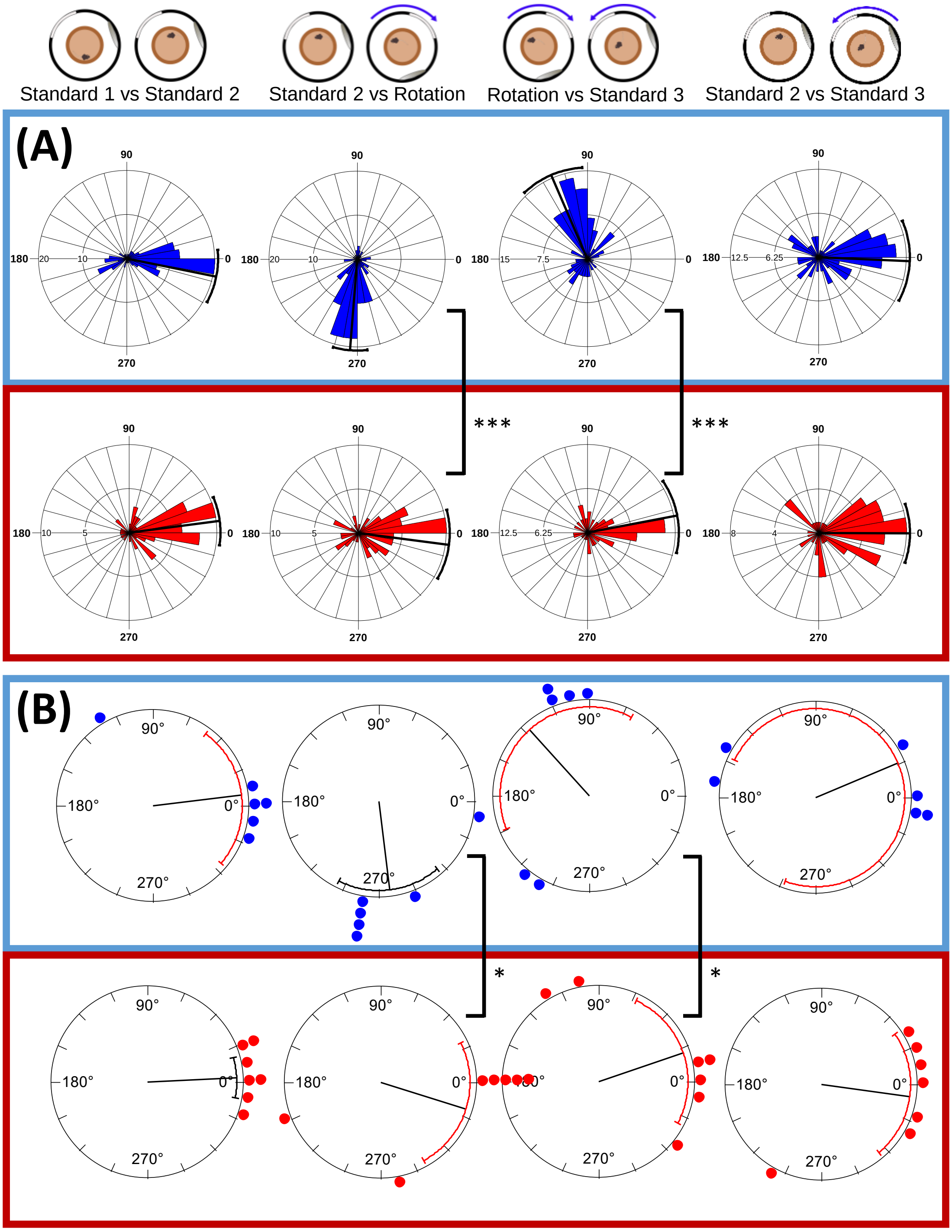


**Figure 2:** Place cells recorded from MEC lesion mice on day 1 of the distal landmark sessions do not rotate with distal landmarks. **(A)** Angular shift at which the maximum correlation between the session rate maps occurred, plotted for cells recorded from all mice in the control (blue) and MEC lesion (red) groups. Radial line denotes circular mean, error bars show 95% confidence intervals. **(B)** Circular mean of maximum correlation angles for cells recorded from each mouse (control, n = 6, lesion, n = 7). Radial line represents mean of the animal means, with 95% confidence interval error bars. * $p < 0.05$, ** $p < 0.01$, *** $p < 0.001$

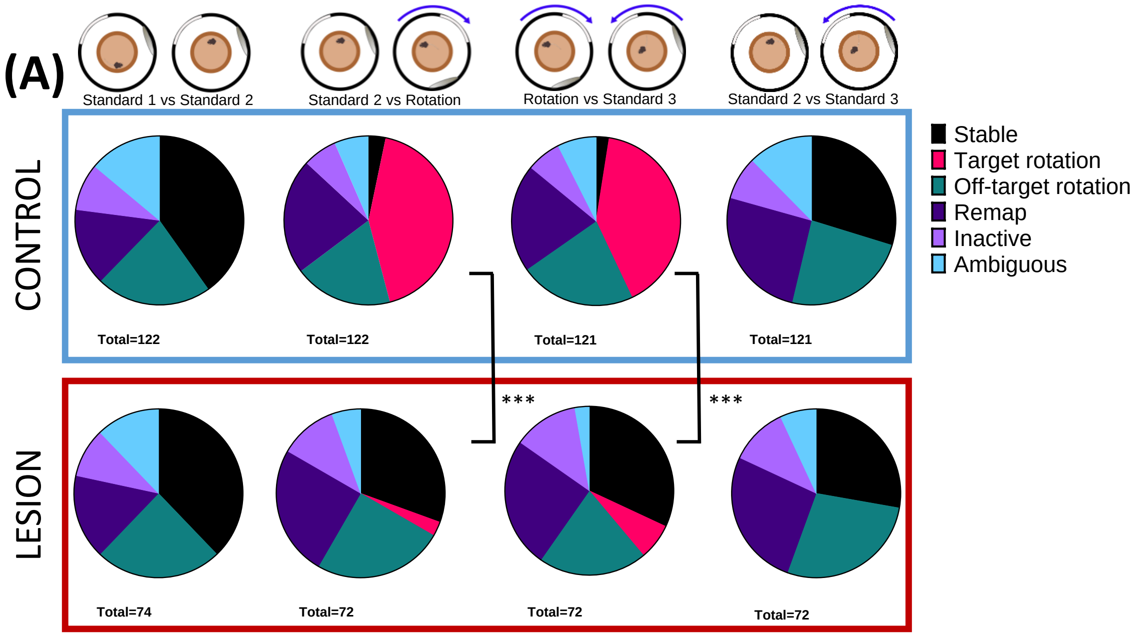


**Figure 3:** Lower proportions of place cells recorded on day 1 rotate with the distal cues in MEC lesion mice. Cells recorded from control (above, blue box) and lesion (below, red box) mice, categorised according to rate map correlations between different sessions, shown as proportions of total number of cells for each session comparison. *** $p < 0.001$

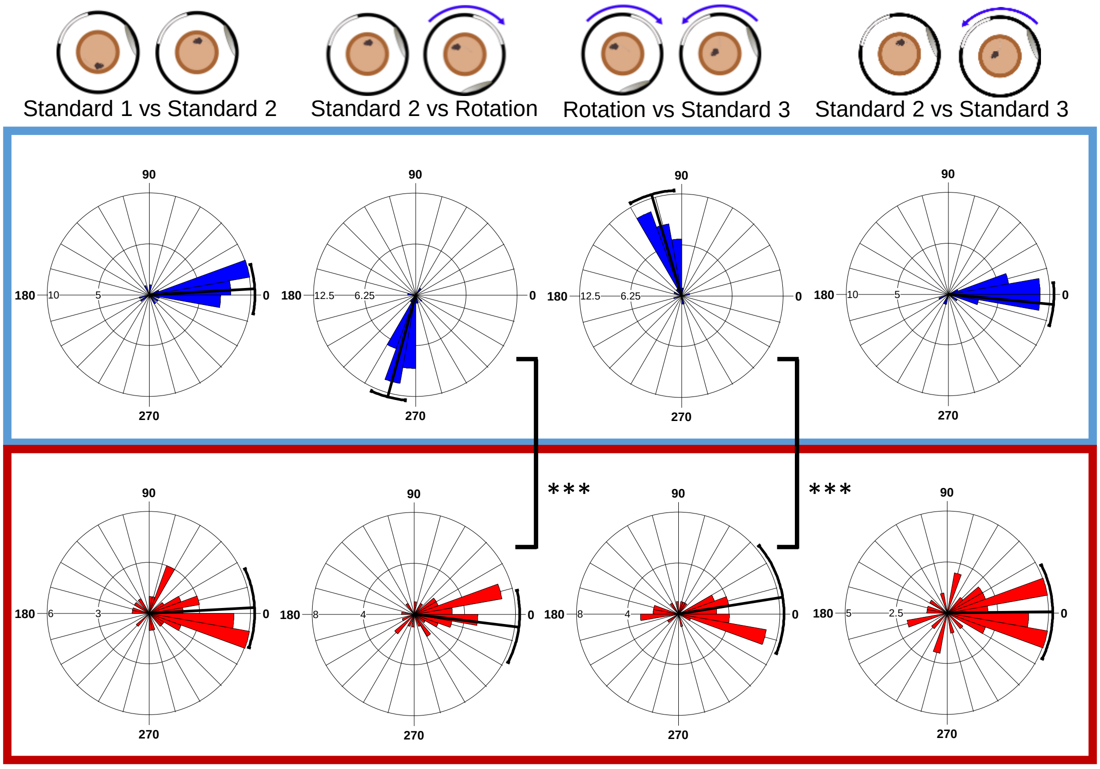


**Figure 4:** Place cells of the control mice show rotations in the distal landmark protocol when recorded following the proximal cue sessions, but cells from lesion mice do not (data collected from 2 control mice and 3 lesion mice). *** $p < 0.0001$

| Mouse # | Lesion type | MEC lesion % | Number of cells | Mean angle | | |
|---|---|---|---|---|---|---|
| | | | | Std1 vs Std 2 | Std2 vs Rot | Rot vs Std3 |
| C1 | Control | 0 | 10 | 1.2 | -47.5 | 105.6 |
| C2 | Control | 0 | 21 | 5.9 | -89.4 | 91.5 |
| C3 | Control | 0 | 73 | -29.2 | -90.3 | 94.5 |
| C4 | Control | 0 | 134 | 24.4 | -93.6 | 42.3 |
| C5 | Control | 0 | 50 | -29.6 | -95.8 | 99.4 |
| C6 | Control | 0 | 36 | 7.0 | -82.8 | 113.0 |
| L1 | Medium | 30 | 13 | 0.3 | -90.8 | 61.4 |
| L2 | Medium | 30 | 11 | 115.8 | -98.7 | 92.2 |
| L3 | Medium | 36 | 24 | 8.1 | -25.0 | 15.2 |
| L4 | Large | 67 | 30 | -7.3 | -9.1 | 25.1 |
| L5 | Large | 71 | 22 | 10.5 | -30.0 | 8.1 |
| L6 | Large | 73 | 54 | 15.4 | -4.4 | 2.3 |
| L7 | Large | 94 | 28 | -0.3 | -7.1 | -19.0 |

**Table 1:** Mean maximum correlation angles of each mouse in the distal cue sessions, with MEC lesion size and number of cells.

| Mouse # | Lesion type | MEC lesion % | Number of cells | Mean angle | | |
|---|---|---|---|---|---|---|
| | | | | Std1 vs Std 2 | Std2 vs Rot | Rot vs Std3 |
| C1 | Control | 0 | 10 | -114.5 | 113.3 | -129.6 |
| C2 | Control | 0 | 21 | -10.5 | -25.3 | 18.1 |
| C3 | Control | 0 | 38 | 1.0 | 46.5 | 68.5 |
| C4 | Control | 0 | 44 | 10.0 | 9.3 | -62.0 |
| C5 | Control | 0 | 40 | 4.5 | -88.2 | 72.2 |
| C6 | Control | 0 | 42 | 1.2 | -73.0 | 79.3 |
| L1 | Medium | 30 | 13 | 47.8 | -140.4 | 101.9 |
| L2 | Medium | 30 | 19 | 18.9 | -76.2 | 76.3 |
| L3 | Medium | 36 | 6 | -43.5 | -80.7 | 42.0 |
| L4 | Large | 67 | 23 | 24.1 | -21.0 | 91.9 |
| L5 | Large | 71 | 23 | -4.1 | -67.7 | 94.1 |
| L6 | Large | 73 | 48 | -7.1 | -72.2 | 71.2 |
| L7 | Large | 94 | 36 | -16.4 | -47.7 | 13.0 |

**Table 2:** Mean maximum correlation angles of each mouse in the proximal cue sessions, with MEC lesion size and number of cells.
